# Genetic predisposition to myeloproliferative neoplasms implicates hematopoietic stem cell biology

**DOI:** 10.1101/790626

**Authors:** Erik L. Bao, Satish K. Nandakumar, Xiaotian Liao, Alexander Bick, Juha Karjalainen, Marcin Tabaka, Olga I. Gan, Aki Havulinna, Tuomo Kiiskinen, Caleb A. Lareau, Aitzkoa L. de Lapuente Portilla, Bo Li, Connor Emdin, Veryan Codd, Christopher P. Nelson, Pradeep Natarajan, Claire Churchhouse, 23andMe Research Team, Björn Nilsson, Peter W.F. Wilson, Kelly Cho, Saiju Pyarajan, J. Michael Gaziano, Nilesh J. Samani, Million Veteran Program, Aviv Regev, Aarno Palotie, Benjamin M. Neale, John E. Dick, Christopher J. O’Donnell, Mark J. Daly, Michael Milyavsky, Sekar Kathiresan, Vijay G. Sankaran

**Author notes:** Correspondence, (V.G.S).

## Abstract

Myeloproliferative neoplasms (MPNs) are blood cancers characterized by excessive production of mature myeloid cells that result from the acquisition of somatic driver mutations in hematopoietic stem cells (HSCs)^1^. While substantial progress has been made to define the causal somatic mutation profile for MPNs^2^, epidemiologic studies indicate a significant heritable component for the disease that is among the highest known for all cancers^3^. However, only a limited set of genetic risk loci have been identified, and the underlying biological mechanisms leading to MPN acquisition remain unexplained. Here, to define the inherited risk profile, we conducted the largest genome-wide association study of MPNs to date (978,913 individuals with 3,224 cases) and identified 14 genome-wide significant loci, as well as a polygenic signature that increases the odds for disease acquisition by nearly 3-fold between the top and median deciles. Interestingly, we find a shared genetic architecture between MPN risk and several hematopoietic traits spanning distinct lineages, as well as an association between increased MPN risk and longer leukocyte telomere length, collectively implicating HSC function and self-renewal. Strikingly, we find a significant enrichment for risk variants mapping to accessible chromatin in HSCs compared with other hematopoietic populations. Finally, gene mapping identifies modulators of HSC biology and targeted variant-to-function analyses suggest likely roles for *CHEK2* and *GFI1B* in altering HSC function to confer disease risk. Overall, we demonstrate the power of human genetic studies to illuminate a previously unappreciated mechanism for MPN risk through modulation of HSC function.

## Main Text

We initially sought to characterize the germline genetic architecture that confers risk for MPNs, and therefore conducted a genome-wide association study (GWAS) meta-analysis using three population-based cohorts (UK Biobank (UKBB), 23andMe, and FinnGen). The combined sample size comprised 2,627 MPN cases and 755,476 controls, more than doubling the number of cases from prior studies (**Supplementary Table 1, Extended Data Fig. 1**). We tested 7,329,649 well-represented variants passing central and study-specific quality control measures. Linkage disequilibrium score regression (LDSC)^4^ showed negligible inflation in test statistics due to population structure, with an intercept of 1.005 and genomic control factor of 1.0255. We estimate the narrow-sense heritability for MPN risk to be 0.0717 (s.e. = 0.0306) on the liability scale, suggesting that ~7% of variance in MPN risk can be attributed to common genetic variation (MAF > 1%), assuming a disease prevalence of 0.0328%, as reported in population studies^5,6^.

Analysis of GWAS signals using GCTA^7^ revealed 12 linkage disequilibrium (LD)-independent loci at genome-wide significance (p < 5 × 10^−8^) and an additional 16 loci with suggestive associations (p < 1 × 10^−6^) (**Fig. 1a, Supplementary Table 2**). Previous studies have shown that carriers of the somatic *JAK2* V617F mutation are strong proxies for MPN cases and can replicate MPN risk associations^8^. Therefore, we assessed genotypes at 27 of 28 lead variants (p < 1 × 10^−6^) among 597 *JAK2* V617F carriers and 220,213 controls in the Million Veteran Program. A near perfect replication for direction of effects was observed (26 of 27 (96%) variants, p = 2.09 × 10^−7^) and 14 variants were significant at the 5% level (p = 6.58 × 10^−12^). We then combined data from MVP with the discovery GWAS to reach a total of 3,224 cases and 975,689 controls. This combined analysis revealed 14 independent risk associations exceeding genome-wide significance, five of which are novel (**Table 1, Supplementary Note**). Of the 10 previously reported loci for MPN risk^8,9^, all but 1 remained genome-wide significant in our analysis, with the only exception being the 22q12.1 locus (p = 9.16 × 10^−7^). We estimate that the 14 genome-wide significant lead variants explain 18.1% of the ~5-fold familial relative risk for MPN acquisition^3^.

**Figure 1.**
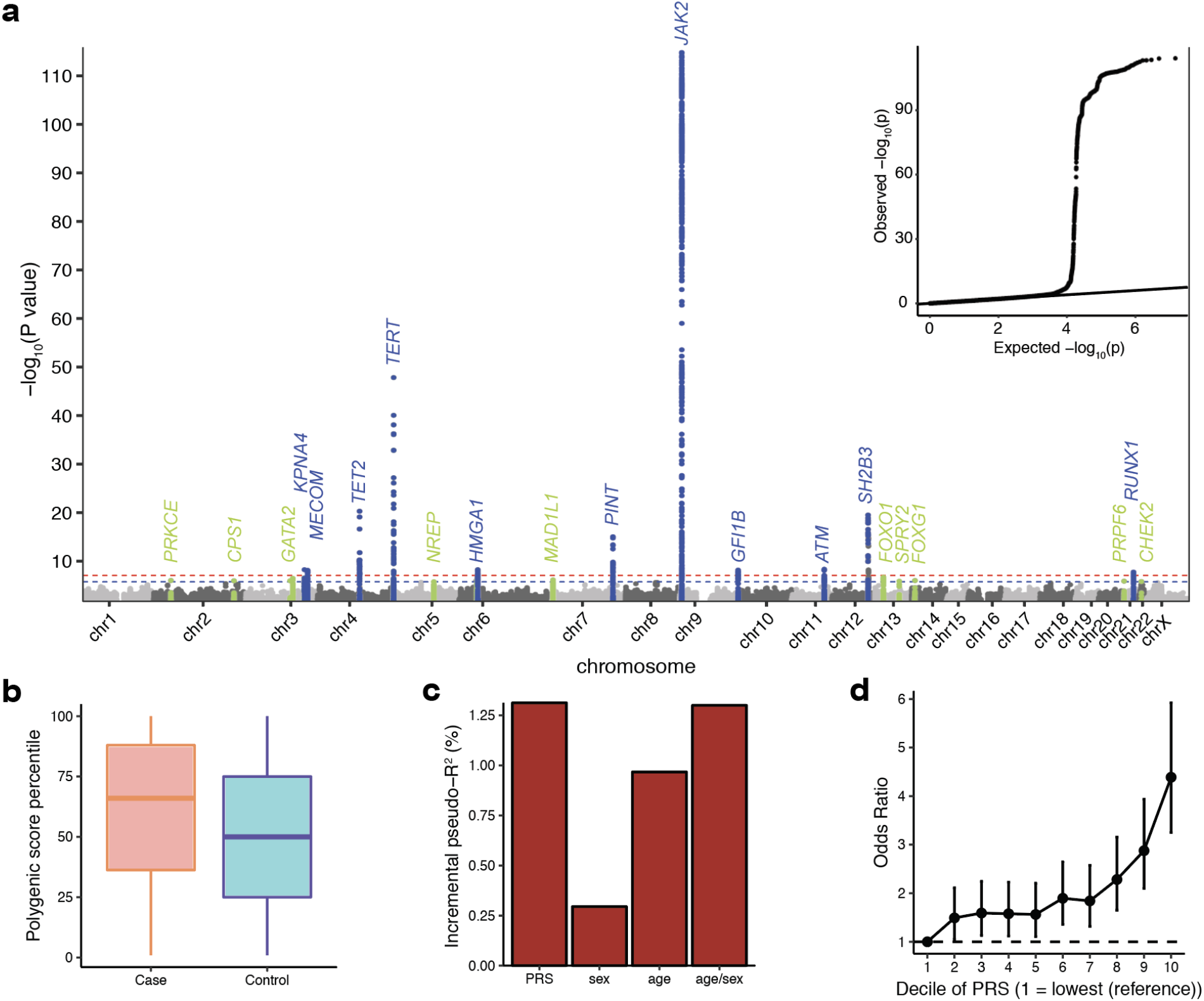
Characterizing the genetic architecture of germline MPN risk. **a**, Manhattan plot and quantile-quantile (QQ) plot (embedded) illustrating results of the genome-wide association study (GWAS) meta-analysis for MPNs in 2,627 cases and 755,476 controls of European descent. The x axis is chromosomal position and the y axis is the significance of association derived by logistic regression. Association signals that reached genome-wide significance (P < 5 × 10^−8^) are shown in blue, and signals that reached suggestive significance (P < 1 × 10^−6^) are shown in green. The QQ plot illustrates the deviation of association test statistics (points) from the distribution expected under the null hypothesis (line). Labels correspond to nearest gene symbols at each distinct association locus (+/- 500 kb). **b**, Polygenic risk score (PRS) percentile among MPN cases versus controls in the UK Biobank testing dataset. Box plots represent the median and interquartile range; whiskers extend 1.5x the interquartile range from the hinges of the box plots. **c**, Additional variance in MPN risk explained by PRS compared to age, sex, and combined age and sex. **d**, Odds ratio (mean and 95% confidence interval) for MPN according to deciles of the PRS, with decile 1 (10% of individuals with lowest PRS) as the reference group.

**Table 1:**
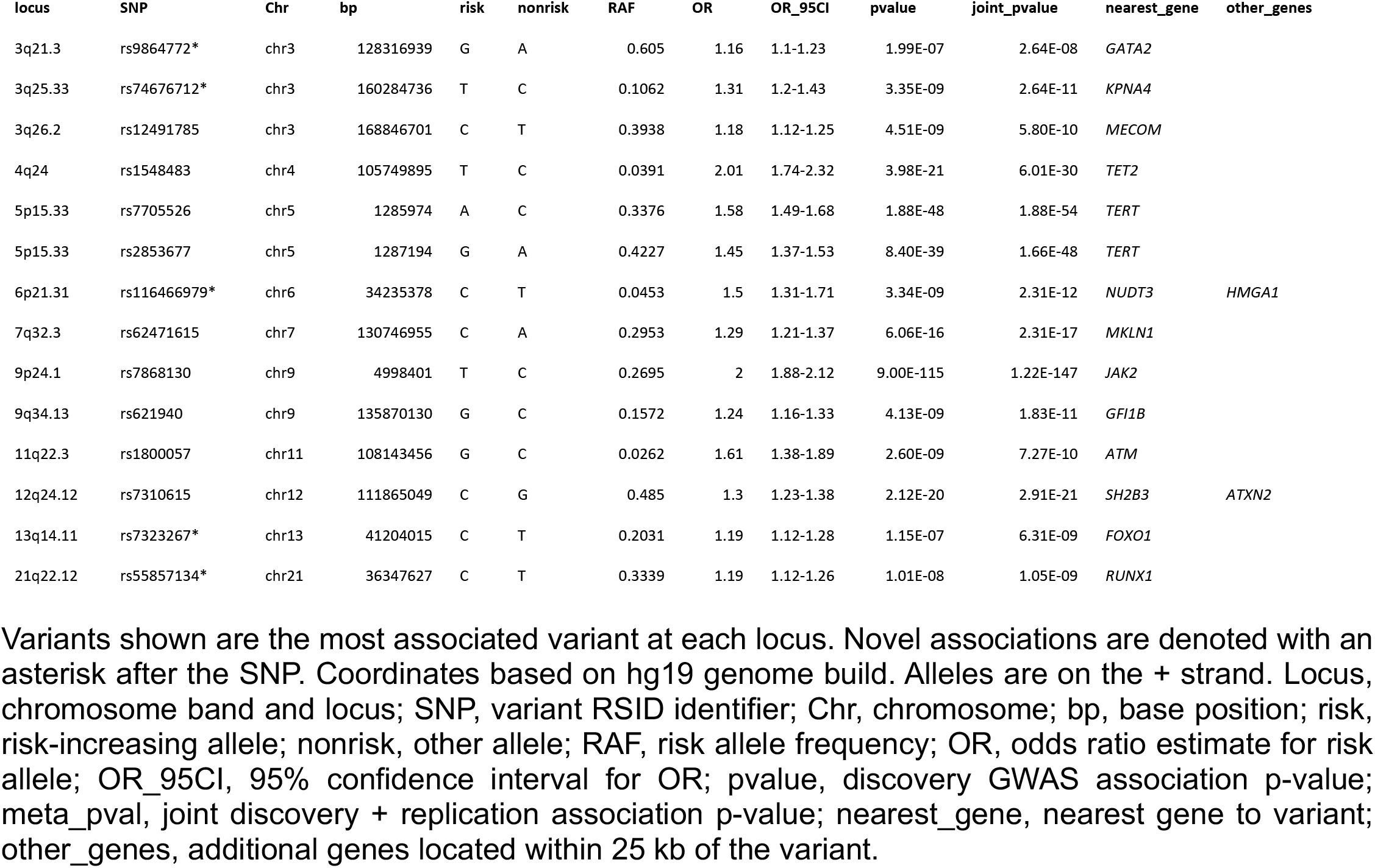
Genome-wide-significant loci from MPN GWAS.

| locus | SNP | Chr | bp | risk | nonrisk | RAF | OR | OR_95CI | pvalue | joint_pvalue | nearest_gene | other_genes |
| --- | --- | --- | --- | --- | --- | --- | --- | --- | --- | --- | --- | --- |
| 3q21.3 | rs9864772* | chr3 | 128316939 | G | A | 0.605 | 1.16 | 1.1-1.23 | 1.99E-07 | 2.64E-08 | *GATA2* |  |
| 3q25.33 | rs74676712* | chr3 | 160284736 | T | C | 0.1062 | 1.31 | 1.2-1.43 | 3.35E-09 | 2.64E-11 | *KPNA4* |  |
| 3q26.2 | rs12491785 | chr3 | 168846701 | C | T | 0.3938 | 1.18 | 1.12-1.25 | 4.51E-09 | 5.80E-10 | *MECOM* |  |
| 4q24 | rs1548483 | chr4 | 105749895 | T | C | 0.0391 | 2.01 | 1.74-2.32 | 3.98E-21 | 6.01E-30 | *TET2* |  |
| 5p15.33 | rs7705526 | chr5 | 1285974 | A | C | 0.3376 | 1.58 | 1.49-1.68 | 1.88E-48 | 1.88E-54 | *TERT* |  |
| 5p15.33 | rs2853677 | chr5 | 1287194 | G | A | 0.4227 | 1.45 | 1.37-1.53 | 8.40E-39 | 1.66E-48 | *TERT* |  |
| 6p21.31 | rs116466979* | chr6 | 34235378 | C | T | 0.0453 | 1.5 | 1.31-1.71 | 3.34E-09 | 2.31E-12 | *NUDT3* | *HMGA1* |
| 7q32.3 | rs62471615 | chr7 | 130746955 | C | A | 0.2953 | 1.29 | 1.21-1.37 | 6.06E-16 | 2.31E-17 | *MKLN1* |  |
| 9p24.1 | rs7868130 | chr9 | 4998401 | T | C | 0.2695 | 2 | 1.88-2.12 | 9.00E-115 | 1.22E-147 | *JAK2* |  |
| 9q34.13 | rs621940 | chr9 | 135870130 | G | C | 0.1572 | 1.24 | 1.16-1.33 | 4.13E-09 | 1.83E-11 | *GFI1B* |  |
| 11q22.3 | rs1800057 | chr11 | 108143456 | G | C | 0.0262 | 1.61 | 1.38-1.89 | 2.60E-09 | 7.27E-10 | *ATM* |  |
| 12q24.12 | rs7310615 | chr12 | 111865049 | C | G | 0.485 | 1.3 | 1.23-1.38 | 2.12E-20 | 2.91E-21 | *SH2B3* | *ATXN2* |
| 13q14.11 | rs7323267* | chr13 | 41204015 | C | T | 0.2031 | 1.19 | 1.12-1.28 | 1.15E-07 | 6.31E-09 | *FOXO1* |  |
| 21q22.12 | rs55857134* | chr21 | 36347627 | C | T | 0.3339 | 1.19 | 1.12-1.26 | 1.01E-08 | 1.05E-09 | *RUNX1* |  |

Variants shown are the most associated variant at each locus. Novel associations are denoted with an asterisk after the SNP. Coordinates based on hg19 genome build. Alleles are on the + strand. Locus, chromosome band and locus; SNP, variant RSID identifier; Chr, chromosome; bp, base position; risk, risk-increasing allele; nonrisk, other allele; RAF, risk allele frequency; OR, odds ratio estimate for risk allele; OR\_95CI, 95% confidence interval for OR; pvalue, discovery GWAS association p-value; meta\_pval, joint discovery + replication association p-value; nearest\_gene, nearest gene to variant; other\_genes, additional genes located within 25 kb of the variant.

To further test the robustness of our findings and their ability to predict MPN risk, we trained a polygenic risk score (PRS) using effect sizes of 92 risk variants obtained from a subset meta-analysis (23andMe and FinnGen), and tested this PRS in the out-of-sample UKBB cohort (**Supplementary Table 3**). Individuals in the UKBB with MPN had a median PRS percentile of 66 compared to 50 for those without MPN (**Fig. 1b, Extended Data Fig. 2**), and the PRS explained a greater proportion of risk variance than that explained by sex and age combined (**Fig. 1c**). The ~40,000 individuals above the 90^th^ PRS percentile had a 4.39-fold (95% CI: [3.25, 5.92]) higher odds of MPN compared to those in the lowest decile (**Fig. 1d**), and 2.78 higher odds compared to those with average genetic risk (5^th^ PRS decile). Altogether, these results indicate that the identified MPN risk associations capture a substantial proportion of total inherited MPN risk and can stratify individual risk.

We next sought to use these genetic associations to gain insights into the mechanisms underlying genetic predisposition to MPNs. Motivated by prior studies showing improved functional enrichments through the inclusion of sub-genome-wide significant associations^10^, we included all 28 suggestive loci in downstream functional analyses. We performed Bayesian fine-mapping on these loci to define credible sets of variants that were 95% probable to contain the underlying causal variant at each signal (**Supplementary Table 4**)^11^. Of the 28 credible sets, 6 (21.4%) contained 5 or fewer variants, and in 17 regions (60.7%), the top fine-mapped variant had a PP ≥ 0.25 (**Extended Data Fig. 3**), enabling refined prediction of causal variants in the majority of regions.

Given that MPNs arise in the hematopoietic compartment, to gain insights into relevant cell states, we examined genetic correlations between MPN risk variants and associations for 19 blood cell traits in 408,241 European ancestry individuals from the UKBB^12^. Strikingly, MPN variants demonstrated significant positive genetic correlations with variants for six diverse blood cell counts – red blood cells, platelets, total white blood cells, monocytes, neutrophils, and eosinophils – all of which derive from multi-potential hematopoietic stem and progenitor cells (HSPCs) (**Fig. 2a-b**). We have previously shown that variants that are pleiotropically associated with distinct blood lineages preferentially fall in accessible chromatin of upstream hematopoietic stem and progenitor populations^13^. Here, a significantly higher proportion of well fine-mapped MPN risk variants (PP > 0.10) demonstrated pleiotropic blood trait associations compared to more weakly fine-mapped variants (PP < 0.10) (**Fig. 2c**, 20.8% vs. 0.8%, Fisher’s exact test p = 2.20 × 10^−9^), supporting the concept that MPN risk variants act in early hematopoietic progenitors.

**Figure 2.**
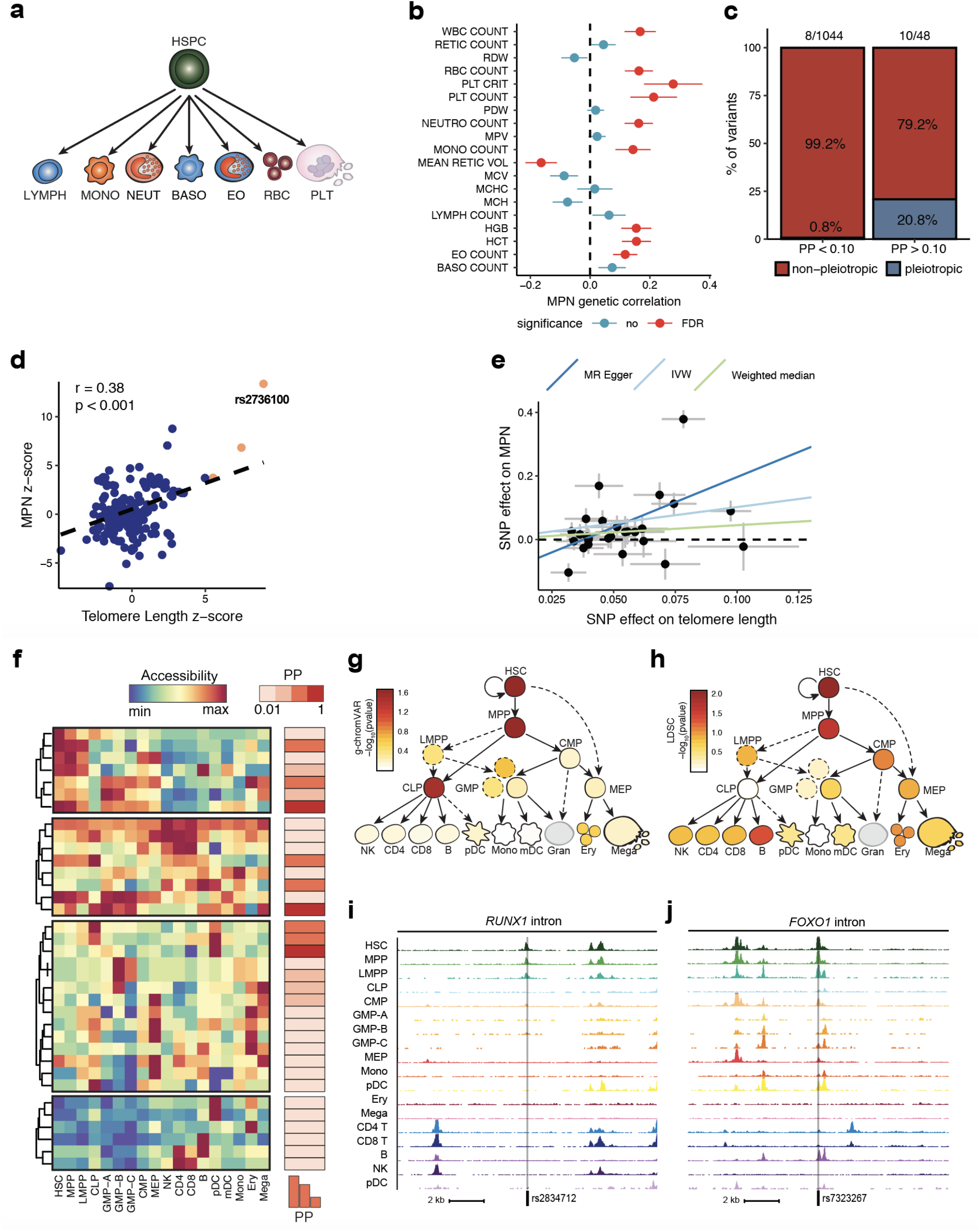
Functional enrichments in MPN risk. **a**, Schematic depicting the trajectory of undifferentiated hematopoietic stem and progenitor cells (HSPCs) into various committed cell types: lymphocytes (LYMPH), monocytes (MONO), neutrophils (NEUT), basophils (BASO), eosinophils (EO), red blood cells (RBC), and platelets (PLT). **b**, Genetic correlations (mean and standard error) between MPN and 19 commonly measured blood traits (results from UK Biobank GWAS of 408,241 individuals): WBC, white blood cell; RETIC, reticulocyte; RDW, red cell distribution width; PDW, platelet distribution width; MCV, mean corpuscular volume; MCHC, mean corpuscular hemoglobin concentration; MCH, mean corpuscular hemoglobin; HGB, hemoglobin; HCT, hematocrit. Red shading indicates significance after adjusting for false discovery rate (FDR-adjusted p < 0.05). **c**, Proportion of fine-mapped MPN risk variants with fine-mapped posterior probability (PP) > 0.10 vs. PP < 0.10 that exhibit pleiotropic associations with blood traits from 2 or more lineages. Absolute proportions are listed above the stacked bar plots. **d**, Z-scores of variants in the *TERT* locus for MPN vs. telomere length. The dashed line represents the linear regression line of best fit. Points colored in orange indicate variants that reach genome-wide significance in both the MPN and telomere length GWAS. **e**, Mendelian randomization scatter plot showing pruned telomere length GWAS variants (p < 1 × 10^−5^) and their effects on MPN risk (outcome) versus telomere length (exposure). Lines represent the slopes of the three methods tested (MR Egger, inverse-variance weighted, weighted median). Error bars represent standard errors of effect sizes. **f**, Heatmap depicting fine-mapped MPN risk variants, clustered by chromatin accessibility across 18 primary hematopoietic populations: B, B cell; CD4, CD4^+^ T cell; CD8, CD8^+^ T cell; CLP, common lymphoid progenitor; CMP, common myeloid progenitor; ery, erythroid; GMP, granulocyte–macrophage progenitor (3 sub-populations); HSC, hematopoietic stem cell; LMPP, lymphoid-primed multipotent progenitor; mDC, myeloid dendritic cell; mega, megakaryocyte; MEP, megakaryocyte–erythroid progenitor; Mono, monocyte; MPP, multipotent progenitor; NK, natural killer cell; pDC, plasmacytoid dendritic cell. Each row marks a fine-mapped variant, each column denotes a hematopoietic cell type, and color denotes relative chromatin accessibility (blue = least accessible chromatin, red = most accessible chromatin). Fine-mapped posterior probability (PP) is indicated to the right. **g-h**, g-chromVAR and LD score regression results for the enrichment of MPN risk variants across chromatin accessibility profiles of 18 hematopoietic cell types. **i-j**, Examples of fine-mapped risk variants with high chromatin accessibility in hematopoietic stem cells (HSCs), located in or nearby genes known to regulate HSC function: (**i**) rs2834712 for *RUNX1* and (**j**) rs7323267 for *FOXO1*.

To gain additional *in vivo* insights into biological connections, we examined the relationship between MPN risk and leukocyte telomere length. Telomere length is associated with HSC self-renewal potential^14^, there is a strong correlation between telomere length of leukocytes and earlier hematopoietic progenitors^15^, and individuals with telomerase loss-of-function associated-diseases have fewer HSCs^16,17^. Motivated by the robust associations for MPN risk at the telomerase reverse transcriptase (*TERT*) locus, we assessed overlap between genetic associations for MPN risk and leukocyte telomere length in 37,684 individuals^18^. At the *TERT* locus, we observed that the lead variant for increased telomere length, rs2736100 (p = 4.38 × 10^−19^), was the second most highly associated variant for increased MPN risk (p = 9.60 × 10^−41^) (**Extended Data Fig. 4**). The top MPN risk variant at this locus, rs7705526, was not genotyped in the telomere length study, but exhibited strong LD with rs2736100 (*r*^2^ = 0.49), suggesting a common signal. Independently, rs7705526 has been found to be associated with clonal hematopoiesis with mosaic chromosomal alterations^19^. In addition, variants in the *TERT* locus common to both MPN and telomere length GWASs demonstrated positively correlated effect sizes (**Fig. 2d**). We then globally tested whether increased telomere length may be causally linked to MPN risk and therefore performed a two-sample Mendelian randomization^20^ using clumped associations from the telomere length GWAS as instruments. Applying four different MR tests with varying assumptions, we consistently found that increased telomere length was significantly associated with increased MPN risk (**Fig. 2e**). The reverse association was also significant using two out of the four tests (see **Methods**). These data show that MPN risk is bi-directionally linked to increased leukocyte telomere length, an important marker of HSC self-renewal capacity. Interestingly, we also noted a suggestive genetic correlation between MPN risk variants and those predisposing to clonal hematopoiesis described in a companion manuscript (r_g_ = 0.39, s.e.m. = 0.21, p = 0.07)^21^, suggesting that these loci may not only promote risk for overt MPNs, but also somatic mutation acquisition in HSCs via a similar mechanism. All of these *in vivo* phenotypic assessments collectively support a role for modulation of HSC function by MPN risk variants.

Given the compelling, albeit indirect, nomination of HSC function by MPN risk variants, we next sought an independent method to holistically assess this concept. We have previously demonstrated the value of using chromatin accessibility (ATAC-seq) data across 18 human hematopoietic cell populations to identify enrichments of fine-mapped variants for hematopoietic traits and thereby enable mechanistic elucidation^13,22^. When we overlapped fine-mapped MPN risk variants with hematopoietic ATAC-seq data, we found that 12.4% (35/282) of MPN risk variants with PP > 0.01 fell within accessible chromatin of one or more hematopoietic populations (**Fig. 2f**), compared to only 5.35% (889/16606) of variants with PP < 0.01 (*χ*^2^ p = 3.95 × 10^−6^), suggesting that stronger fine-mapped variants are enriched for hematopoietic regulatory function. We next used g-chromVAR, a high-resolution cell type enrichment tool^13^, to compute enrichments of fine-mapped variants across these 18 hematopoietic chromatin accessibility profiles. Strikingly, in contrast to common variants associated with blood cell traits, which are maximally enriched in terminally differentiated hematopoietic cells^13^, MPN risk variants showed the strongest enrichment in accessible chromatin of HSCs (p = 2.34 × 10^−2^) (**Fig. 2g**). Because g-chromVAR only considers significantly associated loci (the 28 regions with p < 1 × 10^−6^), we also applied LDSC to assess whether MPN risk is enriched in cell type-specific accessible chromatin across the entire genome. We again found that HSCs showed the highest enrichment relative to all other hematopoietic populations (p = 7.17 × 10^−3^) (**Fig. 2h**). Interestingly, many risk variants locate near known regulators of HSC selfrenewal. For example, rs2834712 (joint p = 1.1 × 10^−9^) lies within an intron of *RUNX1*^23^ (**Fig. 2i**), and rs7323267 (joint p = 6.3 × 10^−9^) lies within an intron of *FOXO1*^24^ (**Fig. 2j**), and both fall in regions with maximal chromatin accessibility in HSCs.

These findings all nominate HSC function as a key target underlying inherited MPN risk. However, we wanted to gain further biological insights into this process by using an integrative approach to map risk variants to target genes. To increase specificity, we focused this analysis on 54 strongly fine-mapped variants with PP > 0.10 and/or lead variants (most significant association) across the 28 suggestive loci. We first nominated three genes from risk loci containing lead variants or variants at PP > 0.10 with missense coding consequences: rs1800057 (risk allele frequency (RAF) = 0.026) causes the P1054R substitution in ATM, rs3184504 (RAF = 0.481) causes the R262W substitution in SH2B3, and rs17879961 (RAF = 0.017) causes the I157T substitution in CHEK2 (**Supplementary Table 5**). For the remaining 25 non-coding risk loci, we prioritized target genes via three independent approaches: 1) if a variant was contained within a gene body, 2) if hematopoietic promoter capture Hi-C (PCHi-C) data^25,26^ suggested that the variant was contained within an enhancer that looped to a promoter, or 3) if there was significant correlation between chromatin accessibility and nearby gene expression across hematopoietic cell types, we would nominate the corresponding gene, as we have successfully done for identifying mechanisms by which blood cell trait-associated variants act^13^. Using these approaches, we identified 28 putative target genes underlying MPN risk. The nominated genes displayed a much stronger set of protein and other functional interactions than expected by chance (STRING database^27^, 18 vs. 7 expected interactions, p = 5.63 × 10^−4^) (**Extended Data Fig. 5**). Remarkably, 10 of the 28 genes have robustly characterized roles as modulators of HSC self-renewal and other functions, including *FOXO1^24^, GATA2^28,29^, RUNX1^23^, PODXL^30^, MECOM^31^, TERT^32^, JAK2^33^, SH2B3^34–36^, ATM^37^*, and *GFI1B^38,39^* (**Fig. 3a**). Consistent with this, the most significantly enriched gene ontology biological processes for the full gene set included replicative senescence, immune system development, and HSC proliferation (**Extended Data Fig. 5, Supplementary Table 6**). Moreover, analysis of bulk RNA sequencing of 16 primary hematopoietic cell populations showed that the 28 target genes are most enriched for expression in HSCs compared to other cell types (rank-sum test, p = 8.7 × 10^−3^) (**Fig. 3a**). To extend these observations to single cell resolution in a more comprehensive survey of human hematopoiesis, we examined the expression of MPN target genes in 278,978 human bone marrow cells. Across 25/28 target genes with detectable expression, we observed a higher MPN score in HSC-enriched cell clusters compared to all other cells. Following gene imputation, we noted a significant co-localization between MPN and HSC signatures across all cells (Spearman rs = 0.27, p < 2.2 × 10^−16^) (**Fig. 3b-c, Extended Data Figs. 6-7**). Together, these results indicate that our nominated set of MPN target genes are enriched for HSC function and expression.

**Figure 3.**
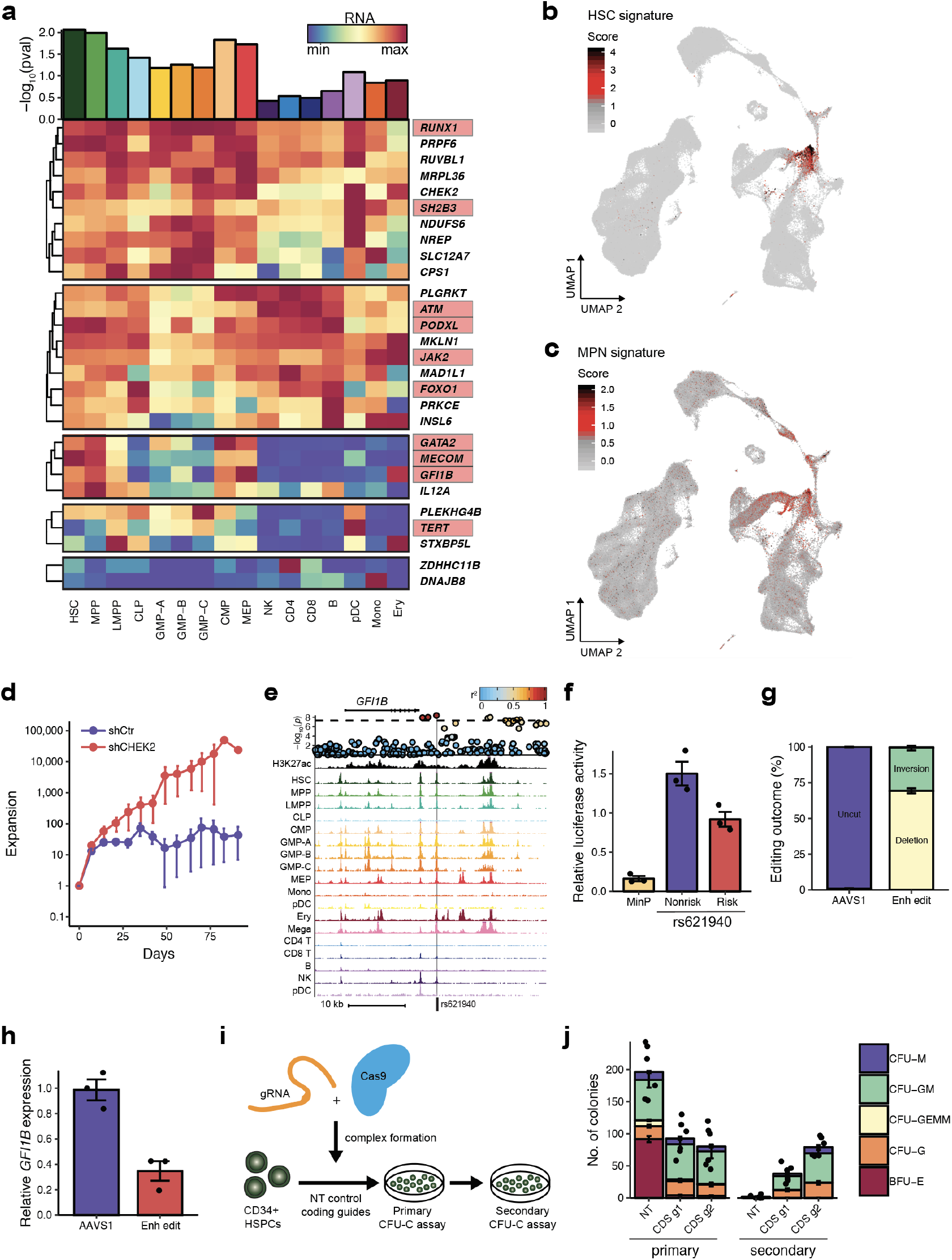
Mapping MPN risk variants to target genes. **a**, Heatmap depicting gene expression across 16 hematopoietic populations for 28 target genes of MPN risk loci. Each row marks a target gene, each column denotes a hematopoietic cell type, and color denotes relative gene expression (blue = lowest expression, red = highest expression). Bar plot above the heatmap depicts the enrichment of target gene expression in each cell type, based on a rank-sum permutation test. Genes which have been previously shown to be involved in hematopoietic stem cell function are boxed in red. **b-c**, UMAP projections of 278,978 single cells from human bone marrow, colored according to (**b**) HSC and (**c**) MPN target gene signatures. d, Expansion of Lin-CD34^+^ derived hematopoietic stem and progenitor cells after short hairpin RNA knockdown of *CHEK2* vs. control. **e**, MPN risk variant rs621940 falls in a region of HSPC H3K27ac signal (top row) and hematopoietic chromatin accessibility (all subsequent rows) ~3 kb downstream of *GFI1B*, a transcription factor known to maintain HSC quiescence. A locus plot showing all MPN GWAS variants in this region is shown above, plotting −log_10_(p) of association; color reflects linkage disequilibrium to lead variant rs621940; the dashed line marks the threshold of genome-wide significance (p = 5 × 10^−8^). **f**, Reporter assays demonstrate allele-specific enhancer activity of rs621940 in hematopoietic cells, compared to a minimal promoter (MinP). g, CRISPR/Cas9 disruption of *GFI1B* super-enhancer (Enh edit) in human CD34^+^ HSPCs results in 99.6% editing rate (69.4% deletion, 30.2% inversion), compared to 0.8% editing rate in a negative control AAVS1 guide. **h**, *GFI1B* super-enhancer disruption in human HSPCs leads to 65% reduction in *GFI1B* gene expression compared to AAVS1 control. **i**, Human HSPCs were electroporated with Cas9 targeting a coding region of *GFI1B* vs. nontargeting (NT) sequence and plated for primary and secondary colony-forming assays. **j**, *GFI1B* coding disruption (CDS) leads to reduced erythroid primary colony formation compared to NT control, but increased secondary colony formation. CFU-M, colony forming unit-macrophage; CFU-GM, granulocyte macrophage; CFU-GEMM, granulocyte erythrocyte macrophage megakaryocyte; CFU-G, granulocyte; BFU-E, burst forming unit-erythroid. In **d, f, g, h**, and **j**, error bars represent standard error of the mean.

The global analyses above strongly nominate modulation of HSC function as a driver of genetic MPN risk. In two select cases, we performed targeted variant-to-function analyses to gain mechanistic insights into disease predisposition, starting with the missense variant in *CHEK2. CHEK2* has been previously shown to modulate genotoxic responses in leukemia, but its role in primary human HSPCs had not been studied^40^. Importantly, the lead risk variant (I157T) in our study has been associated with increased risk for other cancers and shown to be a hypomorphic allele with functionally impaired activation of downstream effectors^41,42^. Knowing this, we utilized previously characterized methods to suppress its activity or expression^40^. Consistent with observations made in leukemia cells, we found that *CHEK2* inhibition reduced genotoxicity upon irradiation of primitive human Lin^-^CD34^+^CD38^-^ cells, as compared to more differentiated progenitors (**Extended Data Fig. 8**). Remarkably, while we did not observe skewed lineage commitment in colony assays by *CHEK2* inhibition (**Extended Data Fig. 9**), suppression of *CHEK2* through RNA interference increased expansion of human cord blood Lin^-^CD34^+^ in long-term cultures in the absence of genotoxic stress (**Fig. 3d**). These results suggest that *CHEK2* may ordinarily constrain HSPC expansion; reduced *CHEK2* function may promote selfrenewal and thereby increase MPN risk. In a second instance, we identified a compelling MPN risk locus ~3 kb downstream of *GFI1B*, falling within a region of active enhancer-associated histone modifications (H3K27ac) and accessible chromatin in human HSPCs (**Fig. 3e**). The risk locus spanned a super-enhancer defined by H3K27ac signal (**Extended Data Fig. 10**), and its chromatin accessibility across cell types significantly correlated with increased *GFI1B* expression (Pearson *r* = 0.620, p = 2.01 × 10^−5^). A reporter assay of the lead risk variant (rs621940) revealed that this regulatory region conferred hematopoietic transcriptional activity, with the risk variant reducing enhancer activity (**Fig. 3f**). Endogenous perturbation of the super-enhancer encompassing this risk locus in human HSPCs resulted in 99.6% editing efficiency and a 65% reduction in *GFI1B* expression (**Fig. 3g-h, Extended Data Fig. 11**). To assess for effects on HSC function with a more homogeneous endogenous alteration, we performed genome editing-mediated perturbation of the *GFI1B* gene itself in human HSPCs and found reduced erythroid differentiation, but greatly increased self-renewal of progenitors as demonstrated through the presence of increased re-plating capacity in methylcellulose colony assays (**Fig. 3i-j, Extended Data Fig. 12**). Analogous to the results obtained for *CHEK2*, these results suggest that this MPN risk locus reduces *GFI1B* expression in HSPCs and may thereby promote self-renewal.

In summary, we have characterized the germline genetic architecture of MPN risk through the largest GWAS to date and implicate HSC function in underlying this risk through a number of functional analyses. Our discoveries of common genetic variation underlying MPN risk complement other recent studies identifying inherited rare variants that promote clonal hematopoiesis^43^. Understanding the fundamental mechanisms of MPN predisposition may inform the development of novel preventive interventions for the disease, analogous to the implementation of human papillomavirus testing and colonoscopic surveillance to reduce cervical and colorectal cancer incidence, respectively^44,45^. In this context, modulation of HSC function or improved surveillance of this compartment may enable therapies to prevent progression to clonal hematopoietic disorders like MPNs^46–48^. More broadly, our findings may serve as a paradigm for dissecting the underlying basis of cancer predisposition alleles.

## Methods

### UK Biobank GWAS

We performed a GWAS on MPNs using the UK Biobank (UKBB)^49^. Written consent was obtained for all participants. Variant imputation was performed as previously described using a combined 1000 Genomes Phase 3-UK10K panel and Haplotype Reference Consortium (http://biobank.ctsu.ox.ac.uk/crystal/label.cgi?id=263)^49^. The following sample-level quality control filters (previously generated by UKBB) were applied to subset our samples: self-reported British ancestry with similar genetic ancestry based on a principal components analysis of the genotypes, included in kinship inference, no excess (>10) of putative third-degree relatives inferred from kinship, no outlier in heterozygosity and missing rates, and no putative sex chromosome aneuploidy. 408,241 individuals satisfied all of these criteria and were included for genetic association studies.

We curated a definition for the MPN phenotype within UKBB using the following codes: Polycythemia (ICD10 D45; ICD9 2384), chronic myeloproliferative disease (ICD10 D47.1), essential thrombocythemia (ICD10 D47.3, D75.2), familial erythrocytosis (ICD10 D75.0), osteomyelofibrosis (ICD10 D47.4, D75.81), chronic myeloid leukemia (ICD10 C921, C922, C931; ICD9 2051), and chronic myeloproliferative disease (ICD10 D47.1). Individuals were also classified as cases if they had a self-reported cancer, self-reported illness code, or histology of cancer tumor code for polycythemia vera, essential thrombocythemia, myelofibrosis, chronic myeloid leukemia, or malignant mastocytosis.

We used SAIGE version 0.29.4^50^ to perform the GWAS using a generalized linear mixed model that controls for case-control imbalance. To select for variants used to estimate the genetic relatedness matrix (GRM), genotyped variants were filtered using the following criteria: minor allele frequency (MAF) > 0.01, LD pruned with r^2^ threshold of 0.2, Hardy-Weinberg p-value > 1 × 10^−6^, and genotype missingness < 0.01. Principal components of ancestry were calculated with these variants using PLINK2 (--pca approx). Age, sex, genotyping array, and the top 10 principal components were included as covariates when fitting the logistic mixed model. For association testing, 26,942,478 autosomal and 1,039,234 X chromosome variants (excluding the pseudo-autosomal region) with MAF > 0.0001 and Info > 0.6 were included.

We also performed GWAS on 19 continuous blood traits within the same 408,241 individuals and same Info and MAF-filtered variants from the UKBB. These associations were performed using BOLT-LMM v2.3.2^51^, with the same covariates of age, sex, genotyping array, and top 10 principal components.

### 23andMe GWAS

GWAS summary statistics on MPNs from the cohort collected by the personal genetics company 23andMe, Inc. were obtained from a previous study, whose analysis has been described in-depth elsewhere^8^. Written consent was obtained for all participants. Imputation was performed using the August 2010 release of 1000 Genomes reference haplotypes. The MPN phenotype was defined using participant self-report for the following diseases: PV, ET, PMF, post-PV/ET myelofibrosis [MF], systemic mastocytosis, chronic myelogenous leukemia, chronic eosinophilic leukemia, and hypereosinophilic syndromes. For our meta-analysis, we used the revised phenotype in their study which also included carriers of the somatic *JAK2* V617F mutation. There was a total of 1,223 cases and 252,140 controls in this population. The GWAS was performed using a logistic regression model with covariates of age, gender, and the top five principal components.

### FinnGen GWAS

The GWAS on MPNs in the FinnGen cohort was performed using SAIGE version 0.29.4, modified to handle missing genotypes and complete separation in covariates. The FinnGen dataset comprised of 96,499 individuals of Finnish descent, including 318 cases of MPN and 96,181 controls. Written consent was obtained for all participants. The MPN phenotype was defined by individuals with one or more clinical codes in nationwide hospital discharge or cause-of-death registries for polycythemia vera (ICD10 D45; ICD9 238.4; ICD8 208), chronic myeloproliferative disease (ICD10 D47.1), thrombocythemia (ICD10 D47.3; ICD9 238.7B; ICD8 287.2), myelofibrosis (ICD10 C9.45; ICD9 238.7A; ICD8 209), and chronic myeloid leukemia (ICD10 C92.1; ICD9 205.1). Variant imputation was performed using a reference panel of 3,775 whole-genome sequenced individuals from Finland. The GRM was calculated with 49,811 variants that were imputed in every cohort, and which satisfied the following quality control filters: INFO > 0.95, MAF > 0.05, genotype missingness < 0.05, LD pruned with r^2^ threshold of 0.1. Age, sex, the top 10 principal components, and genotyping batch/cohort were applied as covariates to the SAIGE logistic regression model.

For GWAS variant quality control, variants were excluded based on significant allele frequency (AF) differences between Affymetrix batches (association p < 1 × 10^−20^ between one batch and the rest for more than one batch, or AF difference > 10 percentage points between one batch and the rest), and AF differences to imputation panel (3,775 WGS Finns) (AF difference between batch and panel > 10 percentage points for at least two batches or log2(af_batch/af_panel) > 3 for at least one batch).

### GWAS meta-analysis

We aggregated association summary statistics from the UKBB, 23andMe, and FinnGen GWAS used a fixed effects model with inverse-variance weighting of log(odds ratios), as implemented in the METAL software^52^. We meta-analyzed 7,329,649 variants which had association statistics in at least the two largest cohorts (UKBB and 23andMe). Linkage disequilibrium score regression (LDSC) of the meta-analysis showed an LDSC intercept of 1.005 and genomic control factor of 1.0255, indicating negligible inflation in test statistics due to population structure. Therefore, we did not adjust test statistics using genomic control.

### Million Veteran Program replication

We performed replication of our discovery meta-analysis results in 220,810 individuals of European descent from the Million Veteran Program (MVP). Written consent was obtained for all participants. We attempted to replicate the lead variants at 27 / 28 suggestive loci identified from our discovery analysis which also had genotype information within MVP (**Supplementary Table 2**); the one exception was rs75405916 (p = 7.4 × 10^−7^, RAF = 0.0004), which was not detected in MVP. Within the MVP cohort, MPN cases were defined as individuals carrying substantial *JAK2* V617F mutation burden, determined by specifying a threshold variant mutant allele intensity for rs77375493, the genotyping probe for the V617F mutation. To set the threshold, we utilized the previously reported odds ratio of 2.04 between rs7868130 (a *JAK2* 46/1 haplotype variant) and *JAK2* pV617F, as an estimate of the true association between *JAK2* 46/1 and V617F cases^8^. Setting the rs77375493 allele intensity threshold to achieve this odds ratio resulted in a V617F case prevalence of 0.27% within MVP, which was comparable to the previously reported population prevalence of ~0.2%^8^. The slightly higher point estimate may reflect the fact that the MVP cohort is predominantly male and older than other population studies, both of which have been associated with increased rates of V617F^53^. We also attempted to replicate using an MPN definition including Phecodes and ICD9 codes (polycythemia vera (Phecode 200.1), chronic myeloid leukemia (Phecode 204.22), essential thrombocythemia (ICD9 238.71), and myelofibrosis (Phecode 289.1, ICD9 238.76)). However, this phenotype resulted in a weaker directional concordance and fewer replications at p < 0.05 as compared to using solely the *JAK2* V617F carrier phenotype. Thus, we chose to use the *JAK2* V617F carrier definition for final replication, given the potential presence of spurious phenotype designations in this cohort. Logistic regression for each replication variant was performed using the PLINK2 −glm function, with age, sex, and the top 5 principal components included as covariates. Inverse-variance weighted meta-analysis was used to compute joint p values combining the discovery meta-analysis and MVP replication associations.

### Approximate conditional association analysis

GCTA was used to perform approximate conditional and joint association analyses (COJO) to identify independent MPN risk loci^7^. In brief, this method performs a stepwise model selection (--cojo-slct) to identify all conditionally independent risk signals at a given p-value of association, using GWAS summary statistics and estimated LD from a reference panel. For estimation of LD, we used a reference sample of 6,000 unrelated individuals of white British origin, randomly selected from the UKBB, similar to the approach performed in a previous study^54^. After excluding variants with low imputation quality (INFO < 0.4) or deviation from Hardy-Weinberg equilibrium (p < 1 × 10^−6^), this reference panel included ~36 million variants. The reference panel was converted from BGEN files to hard-called PLINK files using PLINK2. When running GCTA-COJO, we set the threshold p-value to p < 1 × 10^−6^ and the distance for assuming complete linkage equilibrium (--cojo-wind) at 10000 kb (i.e. 10 Mb).

### Polygenic risk score analysis

We trained a polygenic risk score (PRS) on a meta-analysis of the 23andMe and FinnGen cohorts, which included 1,541 cases of MPN and 348,321 controls. We performed LD clumping on these meta-analysis associations using PLINK version 1.90^55^ (--clump) with an r^2^ threshold of 0.2 and 8 different p-value thresholds: p < 1, 5 × 10^−3^, 1 × 10^−3^, 5 × 10^−4^, 1 × 10^−4^, 5 × 10^−5^, 1 × 10^−5^, 1 × 10^−6^, and 5 × 10^−8^.

We applied the PRS to the UKBB, an out-of-sample test set, containing 1,086 cases of MPN and 401,155 controls. The PRS was computed for each individual in the UKBB by multiplying the genotype dosage of each risk allele for each variant by its association estimate betas (log-odds) as a weight. This was performed using the PLINK2 --score function.

For each p-value threshold, we modeled the PRS using a logistic regression with MPN case-control status as the phenotype and PRS, age, sex, top 10 principal components of ancestry, and genotyping array as covariates. We calculated the area under the receiver-operator curve (AUROC) for each threshold and determined that the p < 1 × 10^−5^ threshold provided the maximum AUROC (0.669). The PRS at this threshold featured 92 clumped risk variants and was used for all further analysis. The proportion of variance explained was calculated by using the Nagelkerke’s pseudo-R^2^ metric. We used this metric to calculate the incremental R^2^, which quantifies the gain in R^2^ when the PRS variable is added to a logistic regression of MPN case-control status on a set of baseline controls (sex, age, genotyping array, 10 principal components of ancestry).

To use PRS to stratify individual risk for MPNs, we divided individuals in the UKBB cohort into deciles based on their PRS. For each of deciles 2-10, logistic models were fitted with the same covariates as used above, comparing MPN risk for members of the given decile compared to those in the lowest decile (i.e., decile 1, containing those with the lowest 10% of PRS).

### Genetic fine-mapping

For each distinct association signal, we calculated approximate Bayes’ factors (ABFs)^11^ for all variants within 1-Mb of the lead variant. ABFs were calculated as:

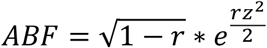

where *r* = 0.04/(*s.e*.^2^+ 0.04) and *z* = *β*/*s.e*. For loci with multiple distinct signals, statistics were based on approximate conditional analysis, adjusting for all other index variants in the region. We then calculated the posterior probability of being causal (PP) by dividing the ABF of each variant by the sum of ABF values over all variants in the locus. The 95% credible set for each locus was constructed by 1) ranking all variants in descending order of PP and 2) including ordered variants until the cumulative PP reached 95%.

### Contribution of variants to overall familial relative risk

We estimated the proportion of the familial risk of MPNs that can be explained by variants identified in our GWAS under a log-additive model, as previously described^56^. We applied the formula 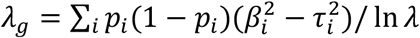, where *λ_g_* is the proportion of familial risk explained, *p_i_* is the MAF for variant *i*, *β_i_* is the log(odds ratio) estimate for variant *i*, *τ_i_* is the standard error of *β_i_*, and *λ* is the overall familial relative risk. We assumed the overall familial relative risk for MPNs to be 4.93 based on a recent epidemiological study^3^.

### Definition of known loci

We compiled a list of 10 previously reported genome-wide significant MPN association signals from the literature^8,9^. Loci were only included if they were identified using a similar MPN phenotype (i.e., combination of *JAK2* V617F carriers and all MPN subtypes, without any sub-stratification of MPN subtypes). Loci which surpassed genome-wide significance (p < 5 × 10^−8^) before or after replication steps were included.

### g-chromVAR cell type enrichment analysis

Bias-corrected enrichment of MPN risk variants for chromatin accessibility of 18 hematopoietic populations was performed using g-chromVAR (https://github.com/caleblareau/gchromVAR), whose methodology has been previously described^13^. In brief, this method weights chromatin peaks by fine-mapped variant posterior probabilities and computes the enrichment for each cell type versus an empirical background matched for GC content and feature intensity. For the chromatin accessibility component, we used a consensus peak set for all 18 hematopoietic cell types with a uniform width of 250 bp centered at the summit. For the variant scores, we used the fine-mapped PP for all MPN risk variants with fine-mapped PP > 0.001 across the 28 suggestive loci.

### Linkage disequilibrium score regression

We used LD score regression (LDSC) to estimate the narrow-sense heritability estimate of MPN risk and compute genetic correlations between MPN risk and other phenotypes^12^. Reference LD scores were computed with a subset of European individuals combined from the 1000 Genomes Phase 3 (1000GP3) and UK10K cohorts. Variants were filtered by MAF > 1%, and 5,653,963 variants were used as input to LDSC. To estimate heritability on the liability scale, the sample prevalence of MPNs was 0.00347 in our GWAS, and the population prevalence of MPNs was estimated to be 0.000328 based on previous reports^5,6^.

To calculate genetic correlations with blood traits, we first used BOLT-LMM to perform GWAS on 19 blood traits in 408,241 European ancestry individuals from the UKBB, the same samples used for the MPN GWAS. Imputation and variant quality control filters were the same as those applied in the MPN GWAS. To calculate cross-trait correlations, we used the same 1000GP3-UK10K reference panel used to estimate LDSC heritability. We constrained the intercept by accounting for the known sample overlap between the MPN and blood trait GWAS, as well as adjusting for phenotypic correlations between MPN case-control status and each of the 19 blood traits. To calculate genetic correlations with blood cancers and cardiovascular disease, we obtained previously generated summary statistics from the UKBB (see URLs) and calculated genetic correlations as described above, except without constraining the intercept.

To calculate cell type enrichments, we generated LD scores for 18 primary hematopoietic ATAC peak sets. We also generated a “pan-heme” peak set representing the union of peaks across all 18 populations. Adopting the approach previously used for LDSC tissue-specific enrichments^57^, we jointly modeled the annotation for each cell type of interest, the “pan-heme” annotation for all hematopoietic peaks, as well as the 52 annotations in the baseline model.

### Blood trait pleiotropy analysis

We tested whether fine-mapped MPN risk variants were more likely to demonstrate pleiotropic associations with common blood traits from distinct lineages. To do this, we performed fine-mapping on all genome-wide significant regions for each of 18 lineage-specific blood traits (all except white blood cell count) using FINEMAP v1.3.1^58^. We then assigned each blood trait to a major hematopoietic lineage: basophil count to basophils; eosinophil count to eosinophils; neutrophil count to neutrophils; red blood cell count, hematocrit, hemoglobin, RDW, MCH, MCHC, MCV, reticulocyte count, and mean reticulocyte volume to red blood cells; platelet count, MPV, platelet crit, platelet distribution width to platelets, monocyte count to monocytes, and lymphocyte count to lymphocytes. We considered a variant to be pleiotropic if it had a fine-mapped PP > 0.10 for blood traits from multiple lineages. Then, we constructed a contingency table for the proportion of MPN fine-mapped variants (PP > 0.10 vs. PP < 0.10), which were classified as pleiotropic vs. not pleiotropic, and calculated an enrichment using a two-sided Fisher’s exact test.

### Mendelian randomization

Mendelian randomization (MR) analysis was performed using the R packages TwoSampleMR^20^ and MRPRESSO^59^. MR consists of two steps: (i) identification of proper instrumental variables or genetic predictors, i.e., variants independently associated with the exposure factor, and (ii) calculation of causal estimates^60^. To achieve step 1, we first obtained GWAS summary statistics for leukocyte telomere length. We then used the UCSC liftOver tool to convert variant coordinates from the hg18 to hg19 genome build. We filtered for variants associated with leukocyte telomere length at a minimum p < 1 × 10^−5^, and then clumped these variants with an LD threshold of *r*^2^ < 0.001 to obtain 27 independent genetic instruments.

Next, we extracted the MPN risk effect sizes of these 27 genetic instruments from our MPN GWAS and harmonized the data to ensure the variant statistics for telomere length and MPN were relative to the same allele. To calculate causal estimates of telomere length on MPN risk, we implemented 4 different methods with varying assumptions regarding horizontal pleiotropy: MR-Egger regression with bootstrap^61^, the IVW method, the weighted median test, and MR-PRESSO. All tests found a significant causal relationship between telomere length and MPN risk: IVW, p = 0.0028; outlier-corrected MR-Presso, p = 0.0015; MR-Egger, p = 0.0031; weighted-median, p = 0.033.

We also tested the reverse association to derive causal estimates of MPN risk on telomere length. The same parameters were used, except now we first clumped the MPN risk variants, and then extracted the corresponding effect sizes for telomere length. The MR test statistics for the reverse association were the following: IVW, p = 0.061; outlier-corrected MR-Presso, p = 0.36; MR-Egger, p = 0.063; weighted-median, p = 0.040.

### Target gene identification

To increase specificity of the target gene analysis, we restricted the input to 54 variants which were either fine-mapped with PP > 0.10 and/or the lead variant at a risk locus (to allow for representation of at least one variant per region). In all analyses, the HLA locus (chromosome 6:28866528-33775446) was excluded due to its complex linkage structure.

We first checked whether any of these variants resulted in coding mutations or splice alterations. To check for splice variants, we annotated all risk variants with spliceAI^62^, a neural net prediction tool for splice altering variants. No variants had a delta score > 0.2, indicating a low probability for any splicing changes. We used Variant Effect Predictor^63^ to screen for coding variants, which identified two variants in distinct loci with PP > 0.10 causing missense mutations: rs1800057 for *ATM* and rs3184504 for *SH2B3*. A lead variant, rs17879961, also caused a missense mutation in *CHEK2*. These three regions were mapped to these respective target genes and were not analyzed further using noncoding variant approaches, described below.

To map the remaining noncoding risk regions to target genes, we incorporated 3 different functional annotations: (1) gene bodies, (2) genes implicated by enhancer-promoter interactions, and (3) chromatin accessibility regions correlated with nearby gene expression. We did not incorporate expression quantitative loci (eQTL) data because to our knowledge, there are no eQTL experiments conducted in hematopoietic bone marrow stem and progenitor cells (HSPCs), the relevant tissue for MPN pathogenesis.

To map variants to gene bodies, we used gene annotation coordinates from GENCODE release 28^64^ for the GRCh37 genome build. We removed all ribosomal protein genes by excluding any genes with names starting with “RP”. We then identified the nearest genes to risk variants using the nearest command in the GenomicRanges R package.

For nominating target genes by enhancer promoter interactions, we used two published promoter capture hi-C datasets^25,26^, spanning 15 terminal hematopoietic cell types and CD34^+^ HSPCs. We filtered for looping interactions with a CHiCAGO score >5. If multiple gene targets were nominated for one variant, only the gene with the top CHiCAGO score was kept.

ATAC-RNA correlations were generated by computing Pearson correlations between hematopoietic ATAC peaks and RNA counts of genes within a 1-Mb window of the ATAC peak, as previously described^13^.

Finally, gene targets from the coding annotations, gene body co-localization, PCHi-C, and ATAC-RNA correlations were combined and filtered for protein-coding genes, as annotated by Ensembl using the annotables R package.

### Target gene cell type enrichments

To measure the enrichment of MPN target genes in bulk RNA-seq data in 16 hematopoietic populations, we performed a rank-sum permutation test. First, we summed the ranks of the 28 target genes ordered by expression (log2 counts per million) amongst all assayed protein-coding genes with non-zero expression in each cell type. Next, we randomly sampled 10,000 equally sized gene sets and obtained their rank sums within each cell type. We calculated the target gene enrichment z-score as the difference between the mean rank-sum of the permuted sets and the target gene rank sum, divided by the standard deviation of the permuted gene set rank-sums. The z-score was then converted into a two-sided p-value for each cell type.

### Gene set enrichments

We used the Functional Mapping and Annotation of Genome-Wide Association Studies (FUMA) tool^65^ to map putative target genes to enriched gene sets. All protein-coding genes were used as the background. Only Gene Ontology Biological Processes were considered.

### Single-cell RNA sequencing analysis

Single-cell RNA-seq of 378,000 cells from human bone marrow were generated as part of the Human Cell Atlas project using 10X Genomics sequencing technology and aligned to the GRCh38 reference genome using the Cell Ranger pipeline as previously described (https://preview.data.humancellatlas.org/). Downstream analyses including normalization, scaling, and cell clustering were performed using the R software package Seurat^66^ version 2 (http://satijalab.org/seurat/). We filtered out low-expressed genes expressed in fewer than 50 cells and low-quality cells with fewer than 500 detected genes, leaving 19,156 genes and 278,978 cells for downstream analysis. Raw gene expression counts of each cell were normalized over total counts and log transformed, and gene expression was then scaled to have a mean of 0 and variance of 1 across cells. We performed dimensionality reduction using PCA, with the top 1000 most variable genes as input, and computed the top 50 principal components (PCs). To identify clusters of cells, we used the ‘FindClusters’ function from Seurat, which applies a shared nearest neighbor modularity optimization-based clustering algorithm to identify clusters based on their PCs (in this case, top 50). To infer the HSC population, we used a marker gene signature similar to one recently applied in a different scRNA-seq human hematopoiesis dataset^67^ – *CD34, HLF*, and *CRHBP*. Of the 28 MPN target genes, we excluded three which were detected in less than 50 cells, resulting in an aggregate MPN signature of 25 genes. To calculate scores based on specific gene sets (e.g., HSC marker genes, MPN target genes) for each cell, we calculated the average of the Z-normalized expression (across all cells) of each gene in the list. To adjust for dropout in single-cell data when estimating the correlation between MPN and HSC signatures, we applied a gene imputation approach called MAGIC to infer missing transcripts in cells. To do so, gene expression value in all cells were normalized, dimensionally reduced and transformed by the internal algorithms in MAGIC with the parameters: n_pca_components = 100, t = 6, k = 10, alpha = 15, rescale_percent = 99. Following imputation, marker gene expression was again used to calculate an MPN and HSC signature per cell, as described above, and a Spearman correlation of these scores was calculated.

### Irradiation experiments

Umbilical cord blood Lineage negative (Lin-) and CD34^+^ cells were obtained using StemSep system according to the manufacturer’s protocol (Stem Cell Technologies, Canada). Lin^-^CD34^+^ cells were sorted to obtain HSC (CD34^+^38^-/low^CD45RA^-^CD90^+^), CMP (CD34^+^38^+^CD45RA^-^CD135^+^), GMP (CD34^+^38^+^CD45RA^+^CD135^+^), and MEP (CD34^+^38^+^CD45RA^-^CD135^-^) fractions. Then cells were resuspended in X-VIVO 10 (BioWhittaker, Waldersville, MD) medium supplemented with 1% BSA, SCF (100 ng/ml), FLT3L (100 ng/ml), TPO (15 ng/ml), G-CSF (10 ng/ml), and IL-6 (10 ng/ml) and incubated for 72-96 hours followed by irradiation with 3Gy. When indicated, cells were pre-treated with CHEK2i (CHEK2 Inhibitor II,10uM final, Sigma 220486) or DMSO for 1hr prior to irradiation. Assessment of IR-induced cell death in the indicated populations 18hr post IR relied on double staining with Annexin and Sytox. IR-induced cell death was calculated to reduce the variability between CD34^+^ batches by subtracting the fraction of AnnexinV+Sytox+ cells scored in the untreated sample from the same fraction in the IR sample.

### Viral constructs, human CD34^+^ transduction and expansion assays

The following pLKO-puro lentiviral shRNA constructs from the RNAi consortium shRNA library (TRC, https://portals.broadinstitute.org/gpp/public/) were used: shCHEK2 (TRCN0000039946) and shControl (TRCN0000231746). This shRNA has been shown to induce knockdown of total and phosphorylated CHEK2 protein^40^. Viral particles pseudotyped with VSV-G were prepared using transient transfection of 293T cells as described elsewhere^68^.

Lin^-^CD34^+^ cells were incubated with the indicated lentiviruses, at multiplicity of infection 50-100, in the X-VIVO 10 medium supplemented with 1% BSA, SCF (100 ng/ml), FLT3L (100 ng/ml), TPO (15 ng/ml), G-CSF (10 ng/ml), and IL-6 (10 ng/ml) for 16 hours followed by cell wash and medium replacement. Puromycin (500ng/ml) was added to the infected cells two days post infection for the additional two days. At the end of puromycin selection CD34^+^ cells were seeded in the ex vivo expansion cultures as previously described^69^. Briefly, CD34^+^ cells were plated in IMDM, 10% FCS (Sigma) supplemented with FLT3L (50 ng/ml), TPO (20 ng/ml), SCF (50 ng/ml), and IL-6 (10 ng/ml) at the density of 1 × 10^5^ cells/ml. Every seven days, cells were counted, washed, and resuspended at the density of 1 × 10^5^ cells/ml in fresh medium and cytokines.

### Super-enhancer analysis

To identify super-enhancers within HSPCs, we utilized previously reported H3K27Ac ChIP-Seq dataset from adult CD34^+^ HSPCs^70^. The raw sequencing data was re-aligned to human genome build hg19 using Bowtie2^71^ with the --very-sensitive parameter, and PCR duplicates were removed with the Picard MarkDuplicates command. ChIP-Seq peaks were called using MACS2^72^ with the --nomodel option and all other parameters left as default. Next, the ranking ordering of super enhancer (ROSE) software^73,74^ was executed on the ChIP-Seq peaks to define super-enhancer status using the following arguments (-s 9000 -t 2500). Enhancers were ranked according to their total reads per million base pairs. An inflection point in the distribution of the occupancy of the factor was used to establish the cutoff for super-enhancers. For visualization, the aligned ChIP-Seq H3K27ac data was converted from BAM to bigWig with bin sizes of 100 base pairs each and normalized to counts per million mapped reads.

### Luciferase reporter assays

The genomic region containing risk and non-risk alleles were synthesized as gblocks (IDT Technologies) and cloned into the Firefly luciferase reporter constructs (pGL4.24) using NheI and EcoRV sites. The Firefly constructs (500ng) were co-transfected with pRL-SV40 Renilla luciferase constructs (50ng) into 100,000 K562 cells using Lipofectamine LTX (Invitrogen) according to manufacturer’s protocols. Cells were harvested after 48 hours and the luciferase activity measured by Dual-Glo Luciferase Assay system (Promega). For each sample, the ratio of firefly to Renilla luminescence was measured and normalized to the minimal promoter construct.

### Ribonucleoprotein electroporation of CD34^+^ human HSPCs

Electroporation was performed using Lonza 4D Nucleofector using 20ul Nucleocuvette Strips. For the large enhancer element deletion experiment, CD34^+^ HSPCs were thawed 24 hrs before electroporation. The RNA complex was made by mixing Cas9 (50 pmol) and modified sgRNAs from Synthego (100 pmol in total), including two guides (sgRNA4: GGCCCAGAAGTGTGGCTGT & sgRNA8: ATGACTTGCTTAGAGCACCA) targeting the five and three prime end of the large enhancer element (hg19 chr9:135868919-135880520). For negative control, a guide targeting AAVS1 site was used (GGGGCCACTAGGGACAGGAT). *GFI1B* gene expression and editing outcome of the electroporation were measured at 6 days post-electroporation. Quantitative PCR was performed using SYBR green (Bio-Rad) to access the frequency of deletion and inversion of the target sequences in bulk cell populations, with primers designed to detect uncut (forward: GAGCCAGCAAAGCCTTAGAA; reverse: GGGAGTATGCAAAGCAGCTC), deletion (forward: TGTCGGTGTCCTGTCTTGAA; reverse: AGACAGCATACGGGGCTAAA), and inversion (forward: GAGTATGCAAAGCAGCTCCC; reverse: CTGCGGGTGGGTTTTCTTAT) events.

For the coding deletion experiment, CD34^+^ HSPCs were thawed 48 hrs before electroporation. The RNP complex was prepared by mixing Cas9 (50 pmol) and modified sgRNA from Synthego (100 pmol) and incubating for 15 min at room temperature immediately before electroporation. HSPCs (3.75 × 10 5) resuspended in 20 μl P3 solution were mixed with RNP and transferred to a cuvette for electroporation with program DZ-100. The electroporated cells were resuspended with Stemspan II media with CC100 cytokine cocktail (Stem Cell Technologies). Two guides targeting *GFI1B* coding regions (sgRNA1: GGGGTCGGGACAGCACAATG; sgRNA2: CCTTGTTGCACTTCACACAG) and control non-targeting guide (NT) was used in these experiments. Gfi-1b protein expression was measured at 5 days post-electroporation.

### Colony-forming unit cell assays

3 days RNP post-electroporation, 500 CD34^+^ HSPCs were plated in 1ml methylcellulose media (# H4034, Stem Cell Technologies). Primary CFU-C colonies were counted after 14 days. For the colony replating experiments, 2 weeks after the primary plating, the colonies from three pates were pooled, washed with PBS, and the cells were plated in new methylcellulose media at 25,000 cells/ml for an additional 2 weeks.

### Data availability

We provide full summary statistics for the MPN meta-analysis comprising UK Biobank and Finngen data at https://www.bloodgenes.org/ (to be uploaded). Summary statistics from analyses based entirely or in part on 23andMe data can only be reported for up to 10,000 SNPs. Thus, all lead variants and the next 8,000 most significant variants from the full GWAS meta-analysis (UK Biobank, Finngen, 23andMe) can be downloaded from https://www.bloodgenes.org/ (to be uploaded). To fully recreate our meta-analysis results for MPN: (1) obtain MPN summary statistics from 23andMe; (2) conduct a meta-analysis of our summary statistics with the 23andMe summary statistics. Individual genetic and phenotypic UK Biobank data are available upon application to the UK Biobank (https://www.ukbiobank.ac.uk).

### Code availability

Code and source data required for reproducing results and figures discussed herein are available on GitHub (https://github.com/sankaranlab/mpn-gwas).

### URLs

g-chromVAR: https://github.com/caleblareau/gchromVAR; Immune cell atlas: https://preview.data.humancellatlas.org/.

## Acknowledgements

We thank members of the Sankaran laboratory for valuable comments. We thank Wei Zhou for technical guidance on the implementation of SAIGE. We would like to thank the research participants and employees of 23andMe. This research has been conducted using the UK Biobank Resource under application 31063. ELB received support from the Howard Hughes Medical Institute Medical Research Fellowship. This work was supported by the Claudia Adams Barr Program for Innovative Cancer Research, National Institute of Health Grant R01 DK103794, and the New York Stem Cell Foundation (VGS). VGS is a New York Stem Cell Foundation-Robertson Investigator.

The following members of the 23andMe Research Team contributed to this study: Michelle Agee, Babak Alipanahi, Adam Auton, Robert K. Bell, Katarzyna Bryc, Sarah L. Elson, Pierre Fontanillas, Nicholas A. Furlotte, David A. Hinds, Karen E. Huber, Aaron Kleinman, Nadia K. Litterman, Jennifer C. McCreight, Matthew H. McIntyre, Joanna L. Mountain, Elizabeth S. Noblin, Carrie A.M. Northover, Steven J. Pitts, J. Fah Sathirapongsasuti, Olga V. Sazonova, Janie F. Shelton, Suyash Shringarpure, Chao Tian, Joyce Y. Tung, Vladimir Vacic, and Catherine H. Wilson.

## Author contributions

E.L.B. and V.G.S. conceived the study. E.L.B, S.K.N., and V.G.S. designed the study. S.K.N., X.L., O.I.G., and M.M. performed experiments. E.L.B., X.L., A.B., J.K., M.T., A.H., T.K., C.A.L., A.L.P., B.L., and C.E. performed compututional and statistical analyses. C.C. and B.M.N. contributed to genetic analysis of UK Biobank. A.B., C.E., P.N., P.W.F.W., K.C., S.P., J.M.G., C.J.O., and S.K. contributed to genetic analysis of the Million Veteran Program. J.K., A.H., T.K., A.P., M.J.D. contributed to genetic analysis of FinnGen. M.T., B.L., and A.R. contributed to analysis of the Human Cell Atlas. V.C., C.P.N., N.J.S. contributed to genetic analysis of leukocyte telomere length. A.L.P., B.N., J.E.D., and M.M. contributed ideas and insights. V.G.S. supervised this work. E.L.B. and V.G.S. obtained funding. E.L.B., S.K.N., and V.G.S. wrote the manuscript with input from all authors. All authors read and approved the final version of the manuscript.

## Corresponding author

Correspondence and requests for materials should be addressed to.

