## Supplementary Figures for "Genetic predisposition to myeloproliferative neoplasms implicates hematopoietic stem cell biology"

### Extended Data Figures

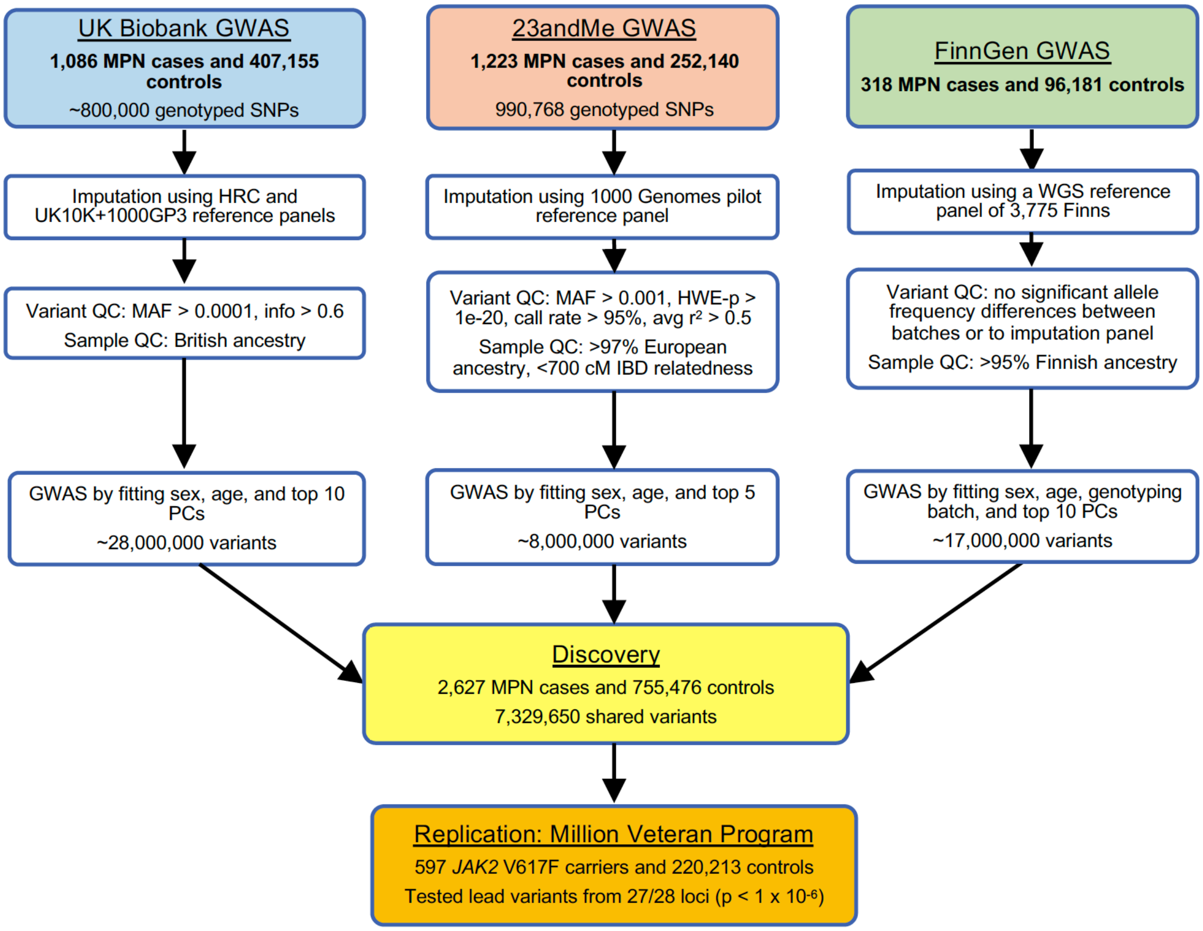


**Extended Data Figure 1.** Flowchart of genetic association analyses. Flowchart of the quality control steps and analysis methods for the three discovery-phase genome-wide association studies (GWAS) in the UK Biobank, 23andMe, and FinnGen, followed by replication in the Million Veteran Program.

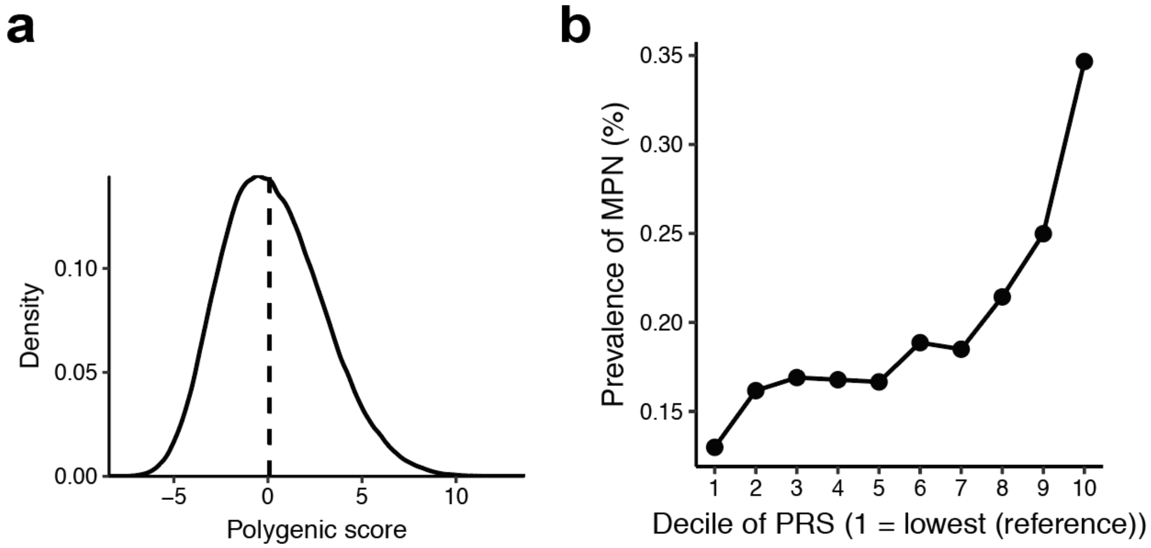


**Extended Data Figure 2.** Assessing the distribution and prevalence of MPN polygenic risk score in UK Biobank. **a,** Density distribution of the MPN polygenic risk score (PRS) within the UK Biobank. **b,** Prevalence of MPN within each decile of the PRS in the UK Biobank population (1,086 MPN cases, 407,155 controls).

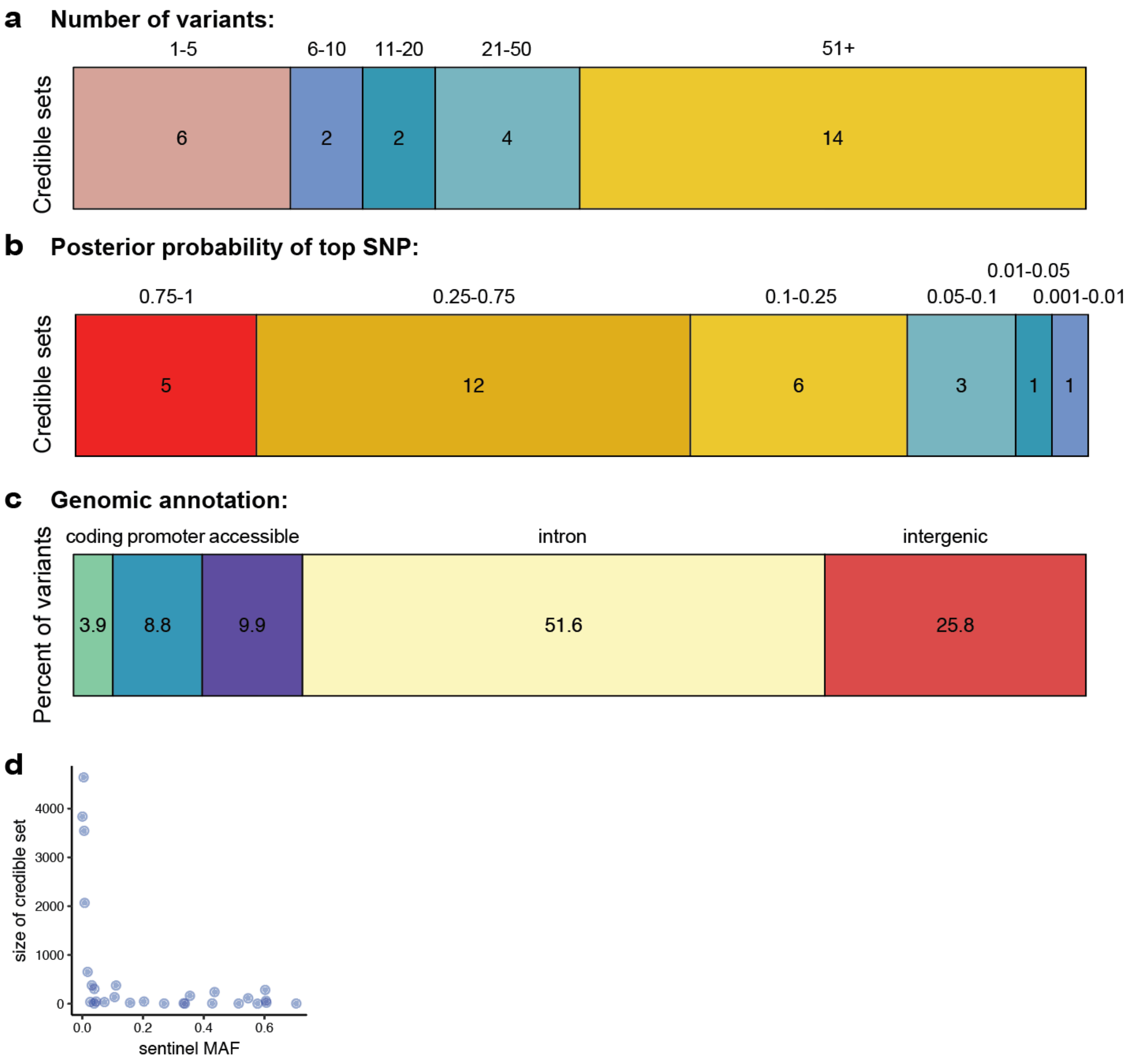


**Extended Data Figure 3.** Descriptive statistics of fine-mapping results for 28 MPN risk loci. **a,** 95% credible sets for all 28 MPN risk loci reaching suggestive significance, stratified by how many variants comprised each credible set. **b,** The fine-mapped posterior probability of causality for the highest fine-mapped variant in each locus credible set. **c,** Variants within the 95% credible sets and posterior probability (PP) > 0.001 across all 28 regions, grouped by genomic annotation. **d,** Size of 95% credible set versus the minor allele frequency (MAF) of the sentinel variant (most significant GWAS p-value) of the credible set.

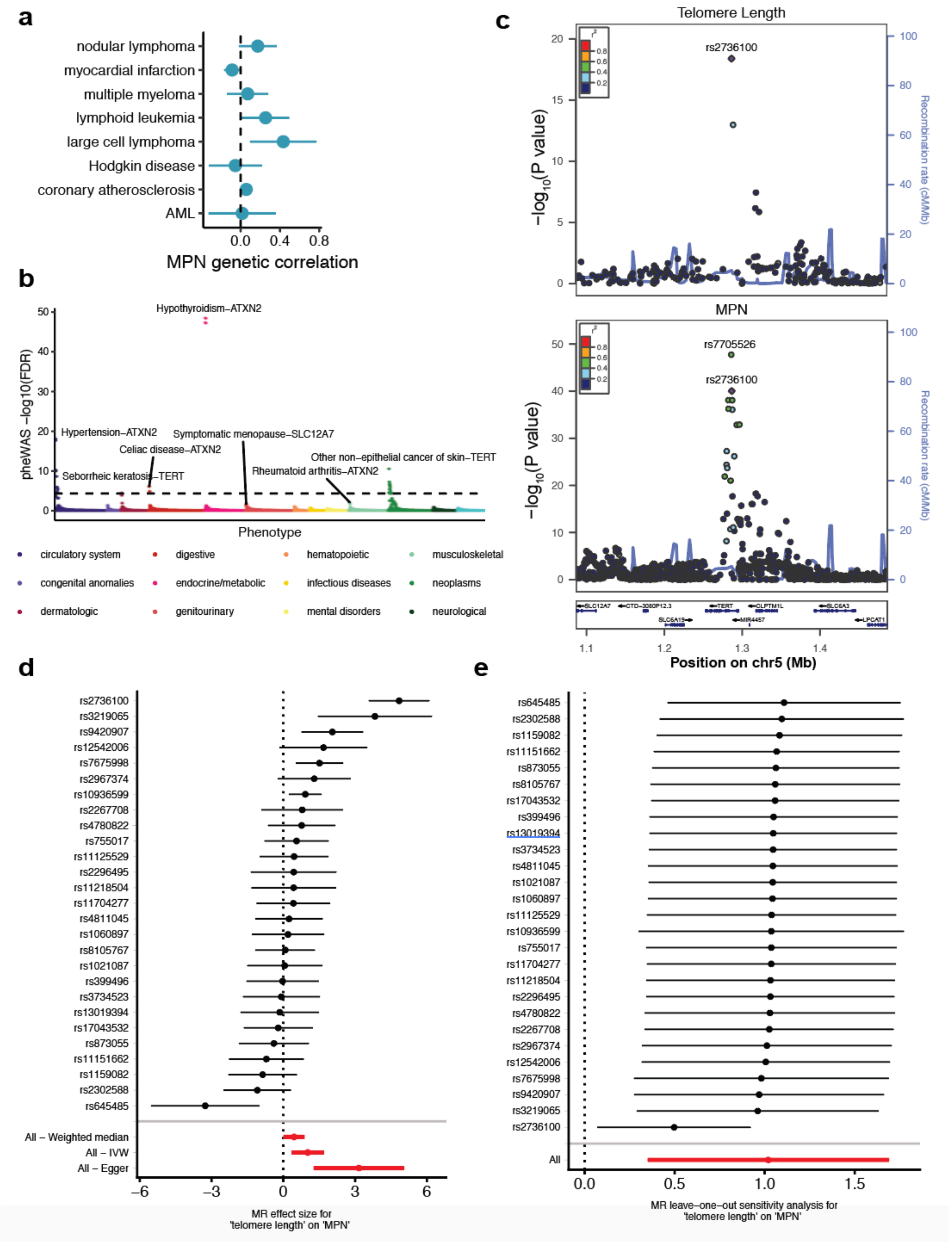


**Extended Data Figure 4.** Associations between MPN risk and other phenotypes. **a,** Genetic correlations (mean and standard error) between MPN risk and other blood cancers and cardiovascular disease, none of which reached statistical significance ($p < 0.05$). **b,** Phenome-wide association study of 51 MPN risk variants (fine-mapped PP > 0.10 or lead variant) for 1,130 clinical phenotypes. Dashed line indicates the FDR-adjusted significance threshold (FDR p = 0.01). The top phenotype association and nearest gene is labeled within each phenotypic category. Full list of significant associations can be found at **Supplementary Table 7**. **c,** Regional association plots at the *TERT* locus, showing the associations of variants with leukocyte telomere length and MPN. In both plots, the colors of the points depict pairwise linkage disequilibrium ($r^2$) to sentinel variant rs2736100. **d,** Individual SNPs associated with telomere length and their effect sizes on MPN risk. Aggregate mendelian randomization (MR) effects, calculated from three different methods (weighted median, inverse-variance weighted, and Egger regression) shown at the bottom. Red color indicates significance. **e,** MR leave-one-out sensitivity analysis, showing the MR effect estimates after excluding each individual SNP from the analysis.

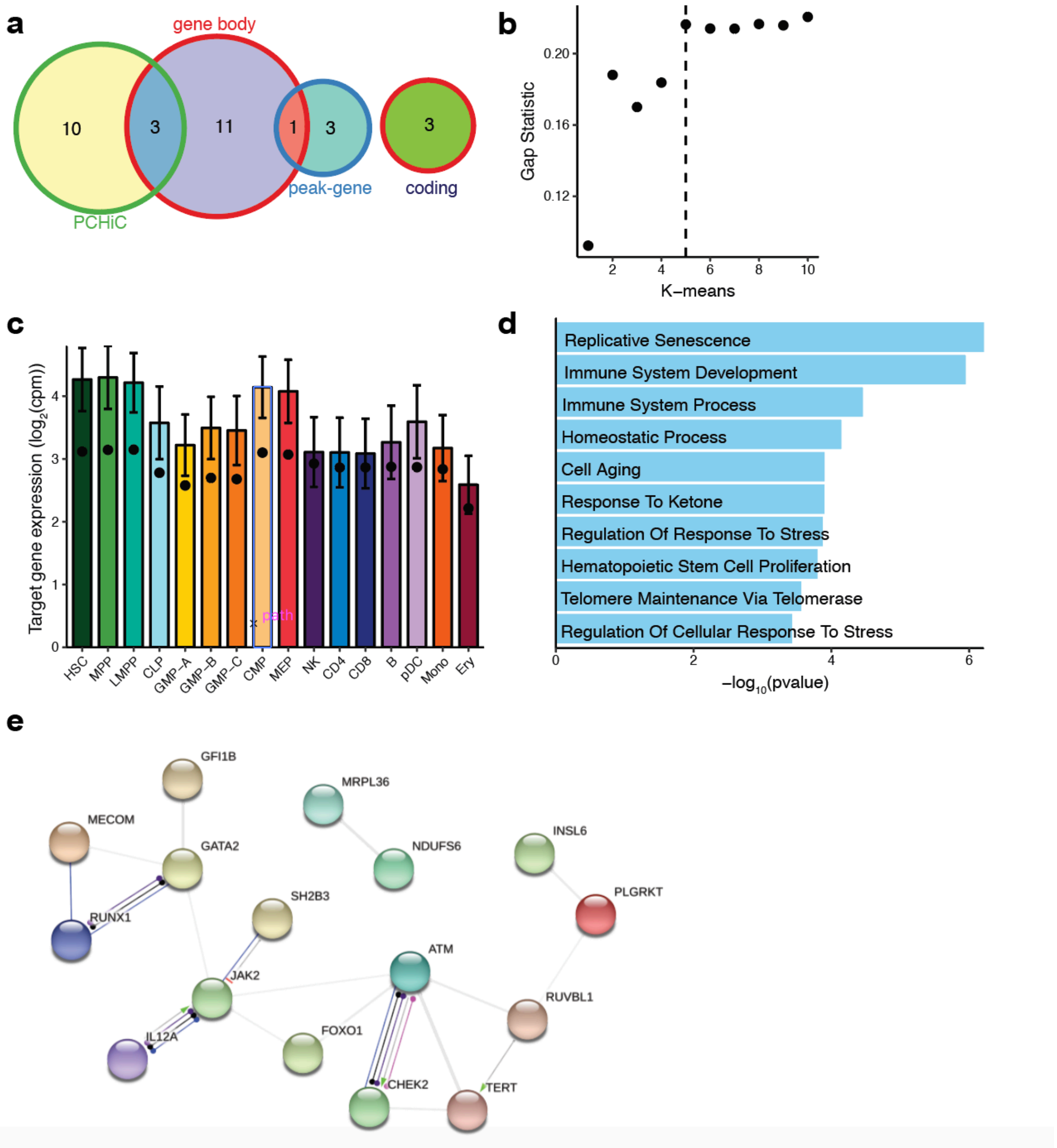


**Extended Data Figure 5.** Characterizing MPN target genes. **a,** Venn diagram showing the intersection of target genes nominated by 1) hematopoietic promoter capture Hi-C (PCHi-C) data, 2) gene body localization, and 3) peak-gene correlations between hematopoietic chromatin accessibility and nearby gene expression. **b,** Gap statistic used to determine number of distinct clusters of target genes, used in **Fig. 3a**. **c,** Expression of MPN target genes (log2 counts per million, mean and standard error) across 16 primary hematopoietic cell types. The dots indicate the mean expression of all non-zero expressed protein-coding genes in each cell type. **d,** Top ten most significantly enriched Gene Ontology biological processes from the MPN target genes.

**e,** Protein-protein interaction network showing known associations between the MPN target genes.

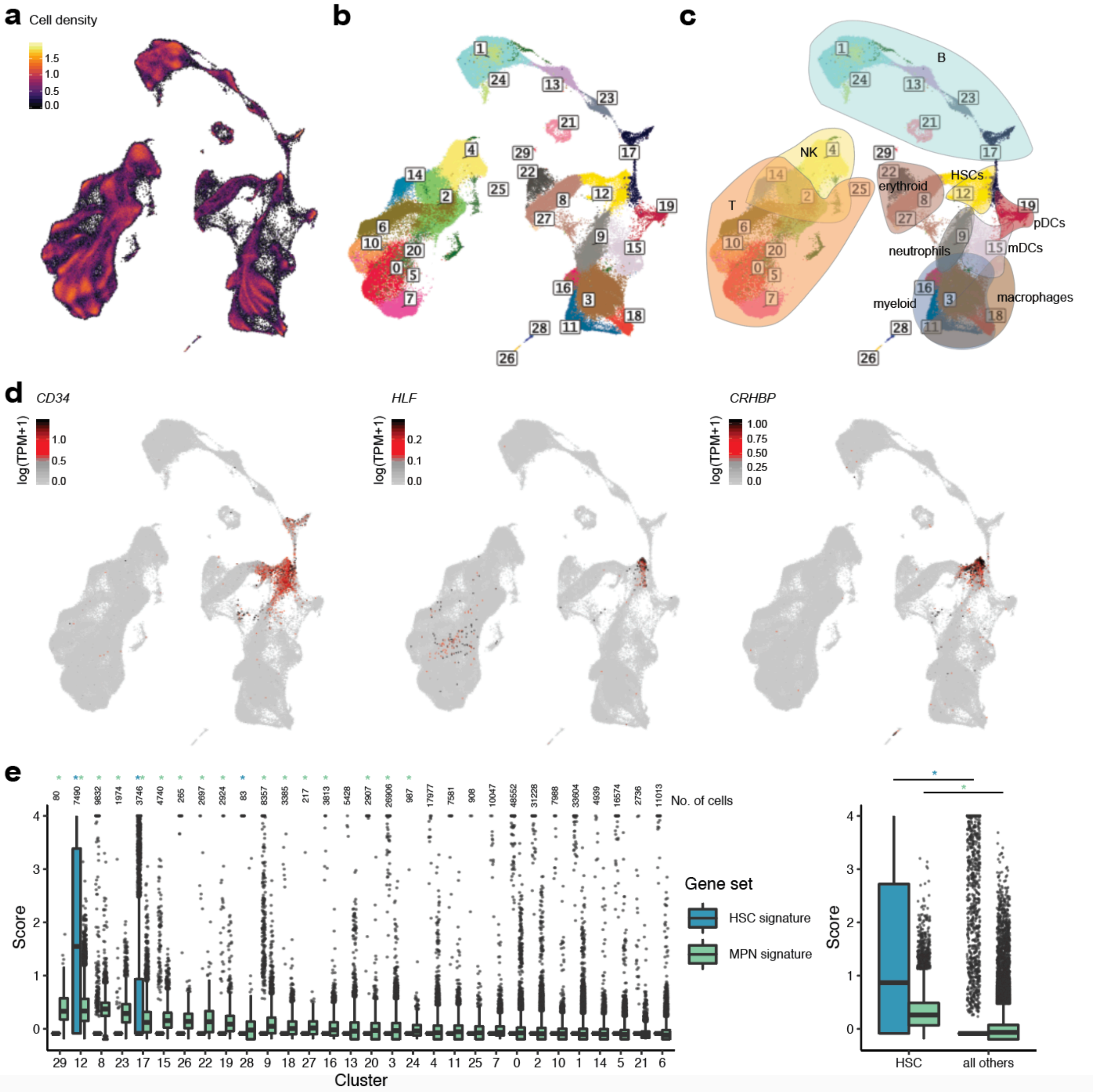


**Extended Data Figure 6.** Inference of cell types within 278,978 single cells from human bone marrow. **a,** UMAP projections of hematopoietic single cells, colored by (**a**) cell density, (**b**) Louvain community clusters, and (**c**) Louvain community clusters with overlays of annotated major hematopoietic lineages inferred from marker genes: B cells marked by *CD79A*, T cells marked by *CD3D*, natural killer (NK) cells marked by *GZMH* and *NKG7*, myeloid cells marked by *FCN1* and *MAFB*, macrophages marked by *CD68* and *CLEC10A*, myeloid dendritic cells (mDCs) marked by *FCER1A*, plasmacytoid dendritic cells (pDCs) marked by IL3RA, neutrophils marked by *ELANE*, and erythroid cells marked by *GYPA*. (**d**) UMAP projections colored by the expression (log(transcripts per million + 1)) of the three gene markers used to annotate HSCs (*CD34*, *HLF*, *CRHBP*). **e,** Left: The distribution of the HSC (blue) and MPN (green) gene scores across all Louvain clusters, ordered from left to right by decreasing average MPN signature score; Right: HSC and MPN gene scores in the combined HSC-significant clusters (12, 17, 28) vs. all other cells. Whiskers extend 1.5x the interquartile range from the hinges of the box plots. False-discovery rate (FDR)-corrected *$P<0.001$ (one-tailed Mann-Whitney U-test), with the * color coded corresponding to the gene signature.

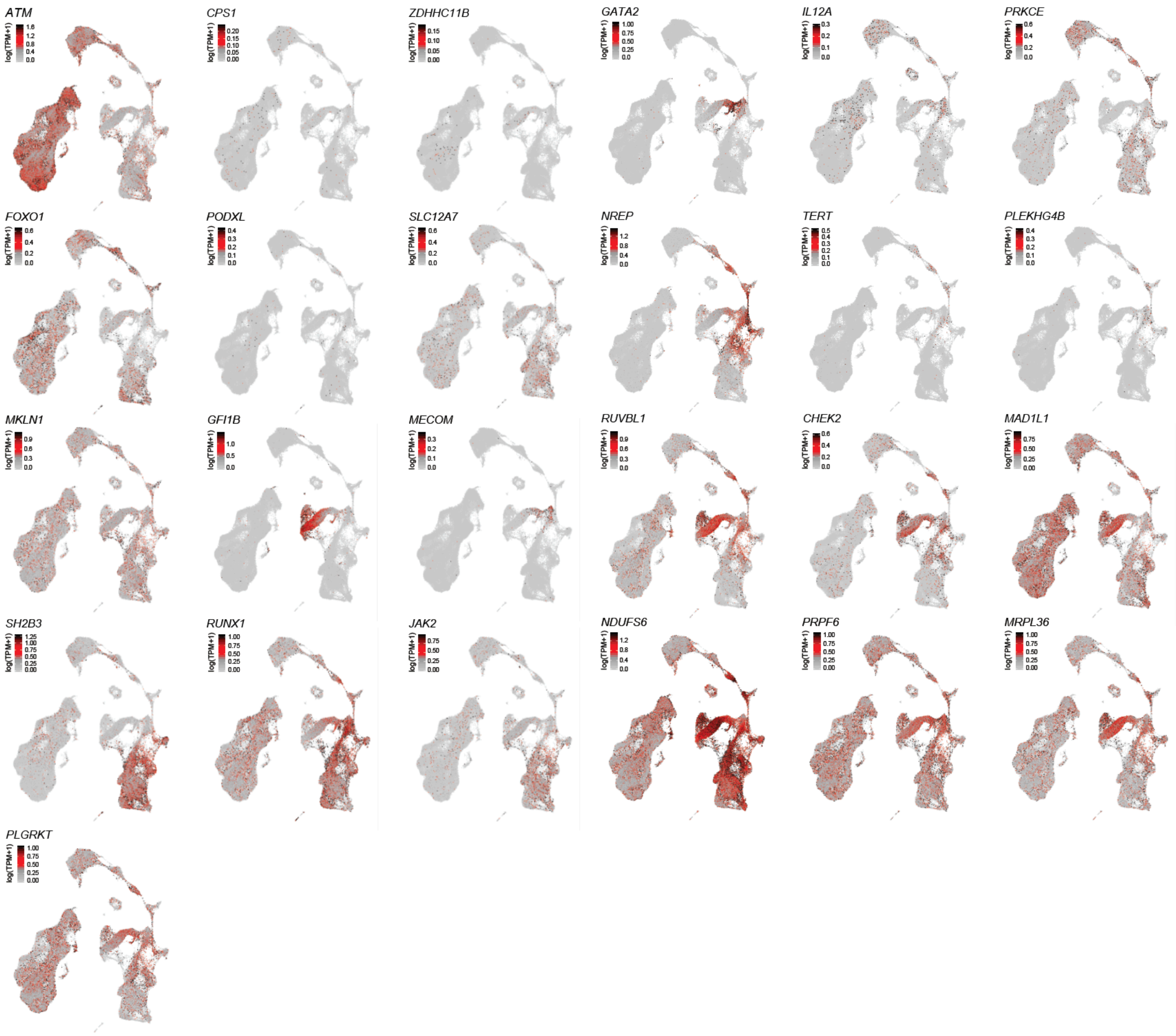


**Extended Data Figure 7.** UMAP projections of hematopoietic single cells, colored by expression of 25 MPN target genes. Color bars represent log(transcripts per million + 1).

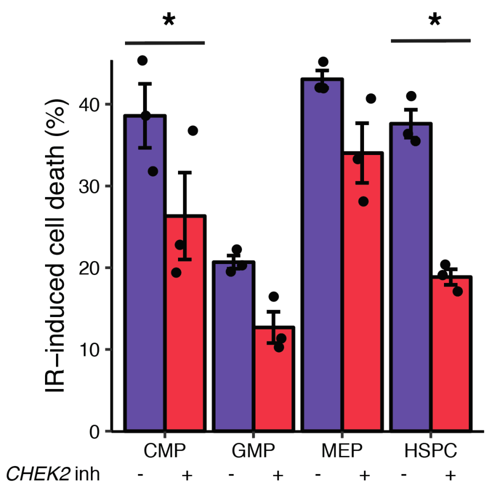


**Extended Data Figure 8.** *CHEK2* is required for apoptosis of cycling HSPCs. Assessment of IR-induced cell death of cycling HSPCs and myeloid progenitors (CMP, common myeloid progenitor; GMP, granulocyte-monocyte progenitor; MEP, megakaryocyte-erythroid progenitor) following sublethal irradiation, after treatment with *CHEK2* inhibitor or dimethylsulfoxide control. *$p < 0.05$, two-sided paired t-test. Error bars denote standard error of the mean.

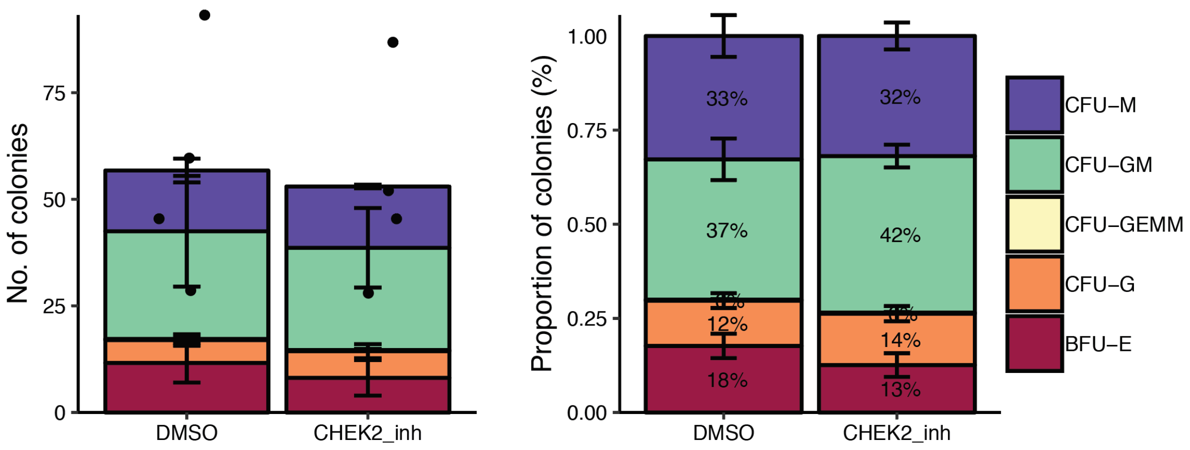


**Extended Data Figure 9.** Numbers (left) and percent (right) of HSPC colonies formed following *CHEK2* inhibition (*CHEK2* inhibitor II, Sigma 220486) vs. dimethylsulfoxide (DMSO) control. CFU-M, colony forming unit-macrophage; CFU-GM, granulocyte macrophage; CFU-GEMM, granulocyte erythrocyte macrophage megakaryocyte; CFU-G, granulocyte; BFU-E, burst forming unit-erythroid. Error bars denote standard error of the mean.

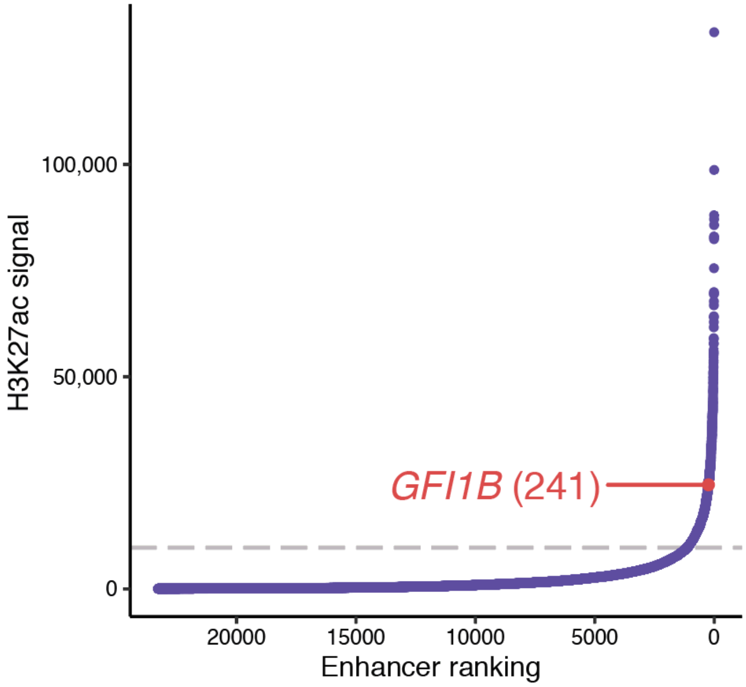


**Extended Data Figure 10.** Distribution of stitched enhancers in adult CD34+ HSPCs, ranked by H3K27ac signal. The dashed line marks the cut-off of 9730.14 used by ROSE to define super-enhancers. The super-enhancer spanning the MPN risk locus near *GFI1B* is ranked 241 and labeled in red.

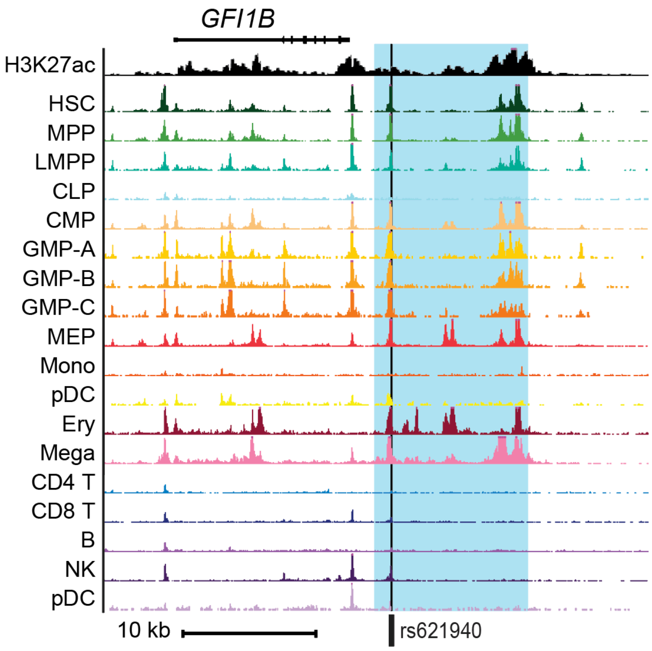


**Extended Data Figure 11.** Genome track depicting the super-enhancer region near *GFI1B* that was targeted for deletion by CRISPR/Cas9 deletion in human CD34$^{+}$ HSPCs (**Fig. 3g-h**). The blue box depicts the targeted deletion, and the black vertical line indicates the lead MPN risk variant at the locus.

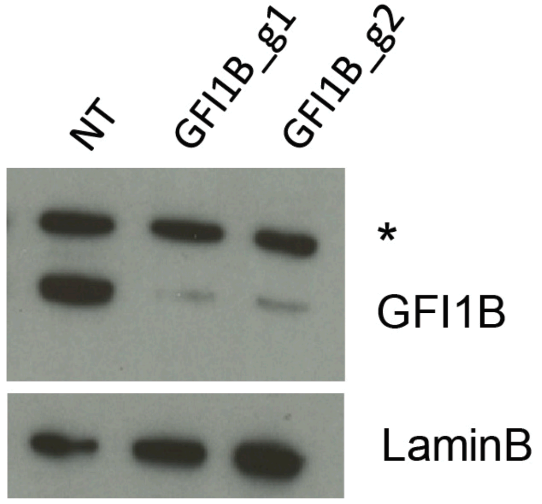


**Extended Data Figure 12.** Western blot of *GFI1B* protein following CRISPR/Cas9 guides targeting non-targeting control (NT), or coding regions of *GFI1B* (g1, g2), compared to LaminB protein expression.
