## Supplementary Note for "Genetic predisposition to myeloproliferative neoplasms implicates hematopoietic stem cell biology"

*Characterizing novel MPN risk loci*

By integrating results from the GWAS, fine-mapping, chromatin accessibility overlap, and gene target analyses, we sought to gain insights into putative biological mechanisms for the five novel genome-wide significant risk loci identified in our study. The majority of association signals involve common non-coding variants, and we found associations near strong candidate genes for MPN risk in pathways or gene families not previously implicated by GWAS.

We identified the first novel genome-wide significant association in locus 3q25.33, represented by lead SNP rs74676712 (RAF = 0.106; p-value = $3.35 \times 10^{-9}$). The nearest gene *KPNA4* encodes importin subunit alpha-4, which has been shown to mediate nuclear localization of STAT3, a downstream mediator of JAK2-mediated signaling[1], as well as other nuclear factors. Importantly, JAK2 itself has been shown to have critical nuclear roles in hematopoietic cells[2], suggesting a potential role for importins in this localization as well.

The second novel locus at 6p21.31, represented by lead SNP rs116466979 (RAF = 0.045; p-value = $3.49 \times 10^{-9}$), is located near *HMGA1*. *HMGA1*, a non-histone chromatin remodeling oncogene, has been shown to be overexpressed in both murine models and patients with polycythemia vera, and higher levels associate disease progression to myelofibrosis and acute myeloid leukemia[3,4]. Interestingly, genetic mutations in the functionally related *HMGA2* gene have also been associated with patients with myeloid neoplasia[5] and hematopoietic stem cell expansion in MPNs[6].

The third novel association was located in locus 21q22.12 within an intron of *RUNX1*. *RUNX1* encodes for runt-related transcription factor 1 and is required for HSC development, HSC homeostasis, lymphoid development, and platelet production. Somatic mutations and chromosomal rearrangements involving *RUNX1* are frequently observed in acute myeloid leukemia[7], myelodysplastic syndrome (MDS)[8], chronic myelomonocytic leukemia[9], and MPNs[10]. Rare germline missense mutations in the gene have been linked to familial platelet disorders with increased risk of myeloid malignancy[11]. Here, we report the first evidence that a common germline variant in *RUNX1* can increase risk of a myeloid malignancy such as MPNs.

The fourth novel region, locus 13q14.11 is represented by the lead SNP rs7323267 (RAF = 0.203; joint p-value = $6.31 \times 10^{-9}$) near the *FOXO1* gene. Previous work has shown that expression of *FOXO1* in human CD34+ cells promotes a preleukemic state with enhanced self-renewal and dysregulated differentiation[12]. Moreover, FoxO1 deletion, in tandem with other FoxO transcription factors, in mice compromises HSC survival[13].

The fifth novel association was found in locus 3q21.3. The lead SNP rs74676712 (RAF = 0.605, joint p-value = $2.64 \times 10^{-}8$) localizes to a distal enhancer for *GATA2*, which encodes a hematopoietic transcription factor that has been shown to play a causal role in inv(3)/t(3;3) AML[14]. Moreover, Gata2 has a critical role in HSC development, self-renewal, and maintenance in mice[15,16]. In addition, human germline mutations in GATA2 compromise HSC function and differentiation[17].

### Supplementary Note References


1 Liu, L., McBride, K. M. & Reich, N. C. STAT3 nuclear import is independent of tyrosine phosphorylation and mediated by importin-α3. *Proceedings of the National Academy of Sciences of the United States of America* **102**, 8150 (2005).

2 Dawson, M. A. *et al.* JAK2 phosphorylates histone H3Y41 and excludes HP1α from chromatin. *Nature* **461**, 819, doi:10.1038/nature08448 https://www.nature.com/articles/nature08448#supplementary-information (2009).

3 Pierantoni, G. M. *et al.* High-mobility group A1 proteins are overexpressed in human leukaemias. *Biochemical Journal* **372**, 145 (2003).

4 Resar, L. *et al.* High Mobility Group A1/2 Chromatin Remodeling Proteins Associate with Polycythemia Vera Transformation to Acute Leukemia in Humans and a JAK2 V617F Transgenic Mouse Model. *Blood* **128**, 1958 (2016).

5 Odero, M. D. *et al.* Disruption and aberrant expression of HMGA2 as a consequence of diverse chromosomal translocations in myeloid malignancies. *Leukemia* **19**, 245, doi:10.1038/sj.leu.2403605 (2004).

6 Ikeda, K., Ogawa, K. & Takeishi, Y. THE ROLE OF HMGA2 IN THE PROLIFERATION AND EXPANSION OF A HEMATOPOIETIC CELL IN MYELOPROLIFERATIVE NEOPLASMS. *FUKUSHIMA JOURNAL OF MEDICAL SCIENCE* **58**, 91-100, doi:10.5387/fms.58.91 (2012).

7 Gaidzik, V. I. *et al.* RUNX1 mutations in acute myeloid leukemia are associated with distinct clinico-pathologic and genetic features. *Leukemia* **30**, 2160, doi:10.1038/leu.2016.126 https://www.nature.com/articles/leu2016126#supplementary-information (2016).

8 Chen, C.-Y. *et al.* RUNX1 gene mutation in primary myelodysplastic syndrome – the mutation can be detected early at diagnosis or acquired during disease progression and is associated with poor outcome. *British Journal of Haematology* **139**, 405-414, doi:10.1111/j.1365-2141.2007.06811.x (2007).

9 Kuo, M. C. *et al.* RUNX1 mutations are frequent in chronic myelomonocytic leukemia and mutations at the C-terminal region might predict acute myeloid leukemia transformation. *Leukemia* **23**, 1426, doi:10.1038/leu.2009.48 https://www.nature.com/articles/leu200948#supplementary-information (2009).

10 Grinfeld, J. *et al.* Classification and Personalized Prognosis in Myeloproliferative Neoplasms. *New England Journal of Medicine* **379**, 1416-1430, doi:10.1056/NEJMoa1716614 (2018).

11 Song, W.-J. *et al.* Haploinsufficiency of CBFA2 causes familial thrombocytopenia with propensity to develop acute myelogenous leukaemia. *Nature Genetics* **23**, 166, doi:10.1038/13793 https://www.nature.com/articles/ng1099_166#supplementary-information (1999).

12 Lin, S. *et al.* A FOXO1-induced oncogenic network defines the AML1-ETO preleukemic program. *Blood* **130**, 1213 (2017).

13 Tothova, Z. & Gilliland, D. G. FoxO Transcription Factors and Stem Cell Homeostasis: Insights from the Hematopoietic System. *Cell Stem Cell* **1**, 140-152, doi:https://doi.org/10.1016/j.stem.2007.07.017 (2007).

14 Gröschel, S. *et al.* A Single Oncogenic Enhancer Rearrangement Causes Concomitant EVI1 and GATA2 Deregulation in Leukemia. *Cell* **157**, 369-381, doi:10.1016/j.cell.2014.02.019 (2014).

15 Rodrigues, N. P. *et al.* Haploinsufficiency of GATA2 perturbs adult hematopoietic stem-cell homeostasis. *Blood* **106**, 477, doi:10.1182/blood-2004-08-2989 (2005).

16 Tsai, F.-Y. *et al.* An early haematopoietic defect in mice lacking the transcription factor GATA-2. *Nature* **371**, 221-226, doi:10.1038/371221a0 (1994).

17 Collin, M., Dickinson, R. & Bigley, V. Haematopoietic and immune defects associated with GATA2 mutation. *British Journal of Haematology* **169**, 173-187, doi:10.1111/bjh.13317 (2015).
